# The *Drosophila* Embryo at Single Cell Transcriptome Resolution

**DOI:** 10.1101/117382

**Authors:** Nikos Karaiskos, Philipp Wahle, Jonathan Alles, Anastasiya Boltengagen, Salah Ayoub, Claudia Kipar, Christine Kocks, Nikolaus Rajewsky, Robert P. Zinzen

## Abstract

*Drosophila* is a premier model system for understanding the molecular mechanisms of development. By the onset of morphogenesis, ~6000 cells express distinct gene combinations according to embryonic position. Despite extensive mRNA *in situ* screens, combinatorial gene expression within individual cells is largely unknown. Therefore, it is difficult to comprehensively identify the coding and non-coding transcripts that drive patterning and to decipher the molecular basis of cellular identity. Here, we single-cell sequence precisely staged embryos, measuring >3100 genes per cell. We produce a ‘transcriptomic blueprint’ of development – a virtual embryo where 3D locations of sequenced cells are confidently identified. Our “*<u>D</u>rosophila*-<u>V</u>irtual-<u>E</u>xpression-e<u>X</u>plorer” performs virtual in situ hybridizations and computes expression gradients. Using DVEX, we predict spatial expression and discover patterned lncRNAs. DEVX is sensitive enough to detect subtle evolutionary changes in expression patterns between *Drosophila* species. We believe DVEX is a prototype for powerful single cell studies in complex tissues.

## INTRODUCTION

Heterogeneous assemblies of cell types acting in concert are the defining feature of complex multicellular life. The metazoan embryo subdivides very early into distinct germ layers that give rise to differentiated cell types, tissues and organs; a process of specialization that requires intricate regulatory networks within and among cells. Cell types differ in gene expression programs that determine fundamental features from morphology to biochemical repertoire, allowing them to fulfill distinct physiological roles. In efforts to better understand the underlying gene expression programs that define cell types, significant efforts have been made to dissect and compare tissue specific materials, either by surgical dissection or genetic and biochemical manipulation. (*1-4*). This has yielded extraordinary insights into, for example, the gene expression dynamics over the course of differentiation and the transcriptomic differences between tissues; however, caveats such as cellular heterogeneity within dissected tissues (*3*) and loss of biological integrity upon genetic manipulation (*4*) have proven difficult to overcome. One approach to avoid heterogeneity is cell culture (*5-11*), but the amount of genomic and transcriptomic variation within most cell lines has been shown to be substantial and cultured cells are often poor *in vivo* proxies.

An alternative is isolation of specific cell types based on expression of markers via cell sorting. This approach has been adapted to genomic studies of complex tissues, including embryos (*12-15*). However, cells are still examined in pools, thus obscuring their significant transcriptomic heterogeneity, which has only recently become appreciable (*16, 17*). The obvious caveat of pooled cell data is that the averaged signatures cannot be de-convolved to the single cell level. For example, specific expression in small subsets of cells may not be detectable, while gene expression relationships such as exclusivity and concomitancy cannot be distilled. This, in turn, limits our ability to infer regulatory mechanisms driving gene expression, as well as to predict the functional roles cells play, how they integrate with, and how they shape their tissue environment.

It is now becoming possible to investigate the transcriptomic landscape of complex cell mixtures with single cell resolution (*18, 19*). Early high throughput single cell expression profiling studies have already allowed for unprecedented insights, from defining the unique relationships between specific cells along differentiation paths, to unveiling lineage branch points, to detecting entirely new cell types in complex mixtures of cells (*17, 20-24*). Single cell transcriptomes thus allow for novel insights into the intricacies of biological specimen and will be essential to reveal the true complexity of even extensively studied model systems like the *Drosophila* embryo.

The *Drosophila melanogaster* (*D. mel.*) embryo has long been an exquisite model for the patterning principles that shape cellular identities. The embryo undergoes 14 rapid nuclear cleavage cycles before cells form during developmental stage 5. The resulting ~6 000 cells in the bilaterally symmetric embryo are morphologically largely equivalent, but within a few minutes cells along the cephalic and ventral furrows invaginate and the germ band extends dorsally (stage 6); cells within the embryo thus begin to act in a coordinated fashion to drive morphogenesis. At stage 5/6, each of the ~3 000 unique cells occupies a specific position along the anteroposterior (AP) and dorsoventral (DV) axes and interprets positional information into transcriptional responses so that any cell’s transcriptomic fingerprint should be a direct corollary of its position within the embryo. Spatially restricted gene expression in early *Drosophila* development has been extensively studied, including efforts to systematically analyze and annotate the expression of a majority of genes (*25-27*). While available *in situ* databases reveal spatial transcription of a host of individual genes, they stop far short of (i) single cell resolution, (ii) allowing the direct comparison of many genes, (iii) covering transcription of the entire genome (including non-coding RNAs), and (iv) allowing for quantitative comparison of gene expression.

Given the relative structural simplicity of the early 6 000 cell embryo and the wealth of gene expression data available, the *Drosophila* embryo should serve as an ideal model for understanding transcriptomic complexity in a differentiating tissue. We show that single cell gene expression profiling of the early embryo allows assaying the transcriptomic state of individual cells genome-wide, quantitatively and without bias. Droplet based sequencing (Drop-seq) enables gene expression profiling of thousands of fixed cells from dissociated embryos at low cost. In fact, *Drosophila* is particularly well suited for Drop-seq, because its small genome allows for sequencing at comparatively high depth and reproducibility. Previously described approaches to spatially resolve single cell transcriptome data (*18, 19*) are not directly applicable for a tissue of our complexity and for the desired spatial resolution; we therefore advanced the mapping strategy by implementing a diffused mapping approach based on Matthews correlation between single cell transcriptomes and a published marker gene set mapped quantitatively and with single cell resolution (*28*). The resulting transcriptomic blueprint of the stage 5/6 *Drosophila* embryo allows us to query the expression patterns of any given gene and map its expression confidently with near-cellular resolution. We verify the predicted expression of known and unknown gene expression patterns, including those of long non-coding RNAs (lncRNAs). We provide an online resource (www.dvex.org) that allows for querying the single cell expression atlas within a virtual embryo gene by gene and combinatorially. The resources provided will not only be useful for the *Drosophila* community, but will allow insights into patterning principles more broadly. The efficacy of such high-resolution transcriptome maps is highlighted in that a comparison of two *Drosophila* species at the mapped single cell level allows for the identification of evolutionary changes in gene expression patterns. This emphasizes the general applicability of single cell transcriptome maps for understanding spatial gene expression dynamics globally, whether the ‘challenges’ are evolutionary time, environmental stress, or genetic aberrations. The mapping and analysis approaches described should be broadly applicable for other systems where complex patterned tissues are to be resolved at the transcriptome level with cellular resolution.

## RESULTS

### Deconstructing the embryo to high quality single cell transcriptomes with Drop-seq

*Drosophila* embryogenesis initiates with the fertilized egg undergoing 14 rapid, largely synchronous nuclear cleavage cycles, resulting in a syncytial embryo of ~6 000 nuclei. By the end of developmental stage 5, the blastoderm embryo has cellularized and spatial gene expression patterns emerge along the AP and DV axes, delineating tissue anlagen, such as the mesoderm (ME) ventrally, the neurectoderm (NE) laterally, and the dorsal ectoderm (DE) dorsally. Stage 6 commences right after cellularization completes, marked by the first morphogenetic movements such as ventral and cephalic furrow invagination and germ band extension. Gene expression around this stage has been assayed in whole embryos (*29, 30*), in maternal effect mutants that convert entire embryos to individual germ layers (*4*), as well as in dissected slices along the AP axis (*31*). These approaches have yielded valuable insights into spatial expression dynamics, but do not allow for cellular resolution. Additionally, removal of an entire body axis by genetic manipulation calls into question the biological identities of the cells and overall integrity of the tissue. On the other hand, large scale *in situ* hybridization projects (*25-27*) capture embryonic expression dynamics of many genes, but this data is neither resolved at the single cell level, nor is it quantitative.

In order to assess wild type transcriptome diversity genome- and embryo-wide at the single cell level, we hand-selected embryos just after onset of gastrulation (stage 6, Fig. 1A). Embryos were dissociated to single cells by dounce homogenization on ice and methanol fixed to ‘freeze’ cells in their developmental state (*32*). Single cell transcriptomes were resolved by droplet-based sequencing (Drop-seq), which employs single cell encapsulation, droplet-contained cell lysis and capture of polyadenylated transcripts by capture probes covalently linked to beads. The capture probes are designed to facilitate first strand synthesis and unambiguous cell assignment (‘stamp’ barcodes) and elimination of library amplification artifacts using unique molecular identifiers (‘UMIs’). We optimized the Drop-seq procedure (*24*) to maximize sequencing quality and minimize doublet rates (Suppl. Methods). Across 7 Drop-seq runs corresponding to 5 biological replicates (Sup. Table S2), >5 000 precisely staged *D. mel.* embryos were dissociated into single cells (Fig. 1A, left, Materials and Methods for details), resulting in a total of ~7 975 sequenced *D. mel.* cells (Suppl. Table S3). To assess doublet rates, 2 of the replicates contained mixtures of cells from *D. mel.* and *Drosophila virilis (D. vir.)* embryos, two species that are separated by > 40 million years of evolution. Accordingly, species specific alignments reveal whether stamp-specific UMIs align to *D. mel., D. vir.,* or both – we observed species doublet rates of ~6.2% and ~3.9% for the two runs (Fig. 2A, Suppl. Tables S2 and S3). Thus, the vast majority (> 90%) of the sequenced cells represent true single cell transcriptomes.

**Figure 1:**
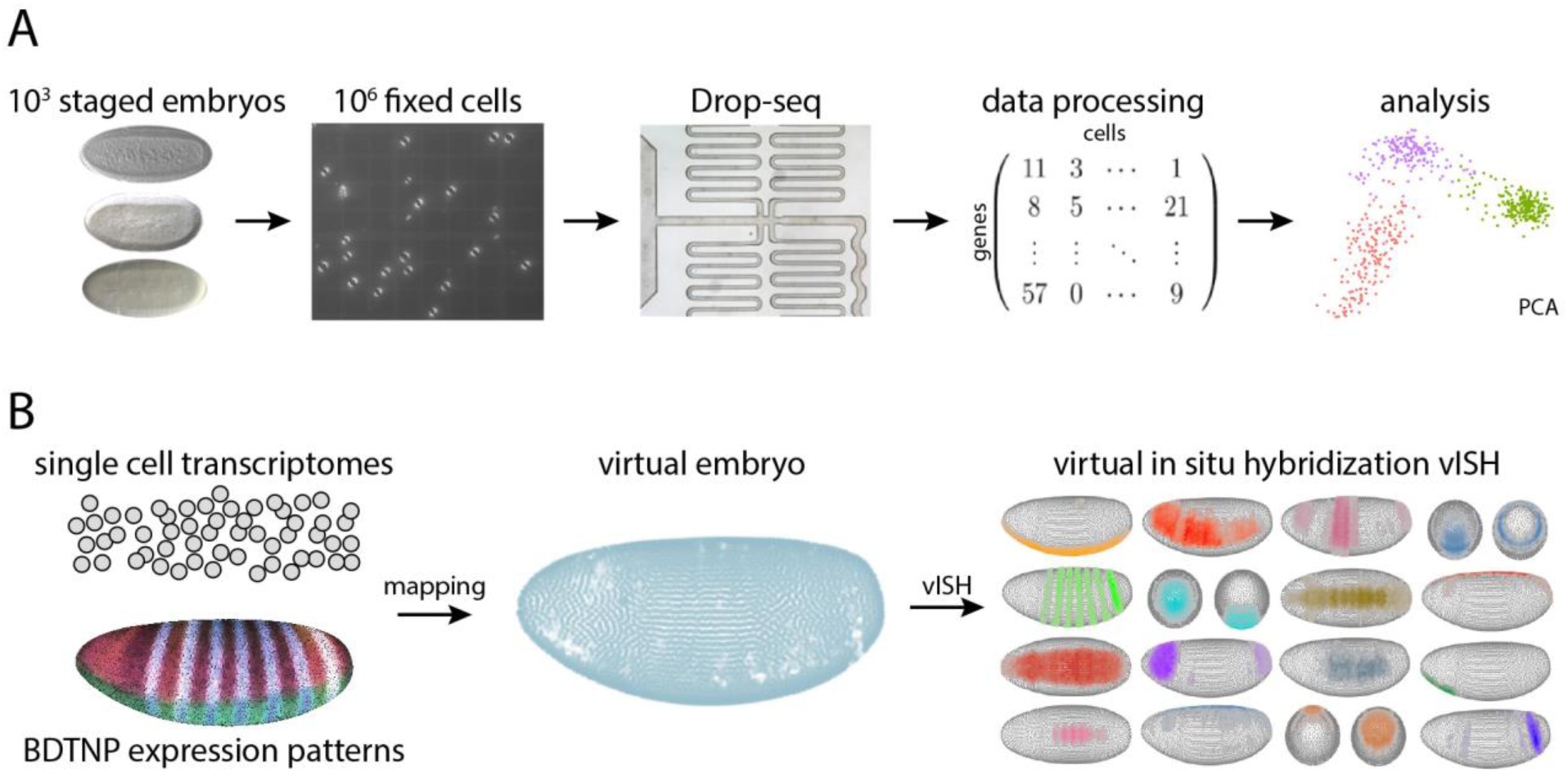
Taking the embryo apart and putting it back together. (A) For each Drop-seq replicate, ~1 000 hand picked stage 6 embryos are dissociated → cells are counted and fixed → single cells are combined with barcoded capture beads in a microfluidic device, followed by library preparation and sequencing. → Single cell transcriptomes are deconvoluted *in silico* resulting in a digital gene expression matrix. The PCA shows separation of cells based on transcriptome, coloring based on expression of marker genes (ME, mesoderm, green; NE, neurogenic ectoderm, purple; DE, dorsal ectoderm, red). (B) The transcriptomes of sequenced cells are combined with high resolution gene expression patterns (*28*) across 84 marker genes, → which allows positional mapping to a virtual embryo. → Virtual in situ hybridization (vISH) predicts which cells express any given gene and where they map within the embryo.

**Figure 2.**
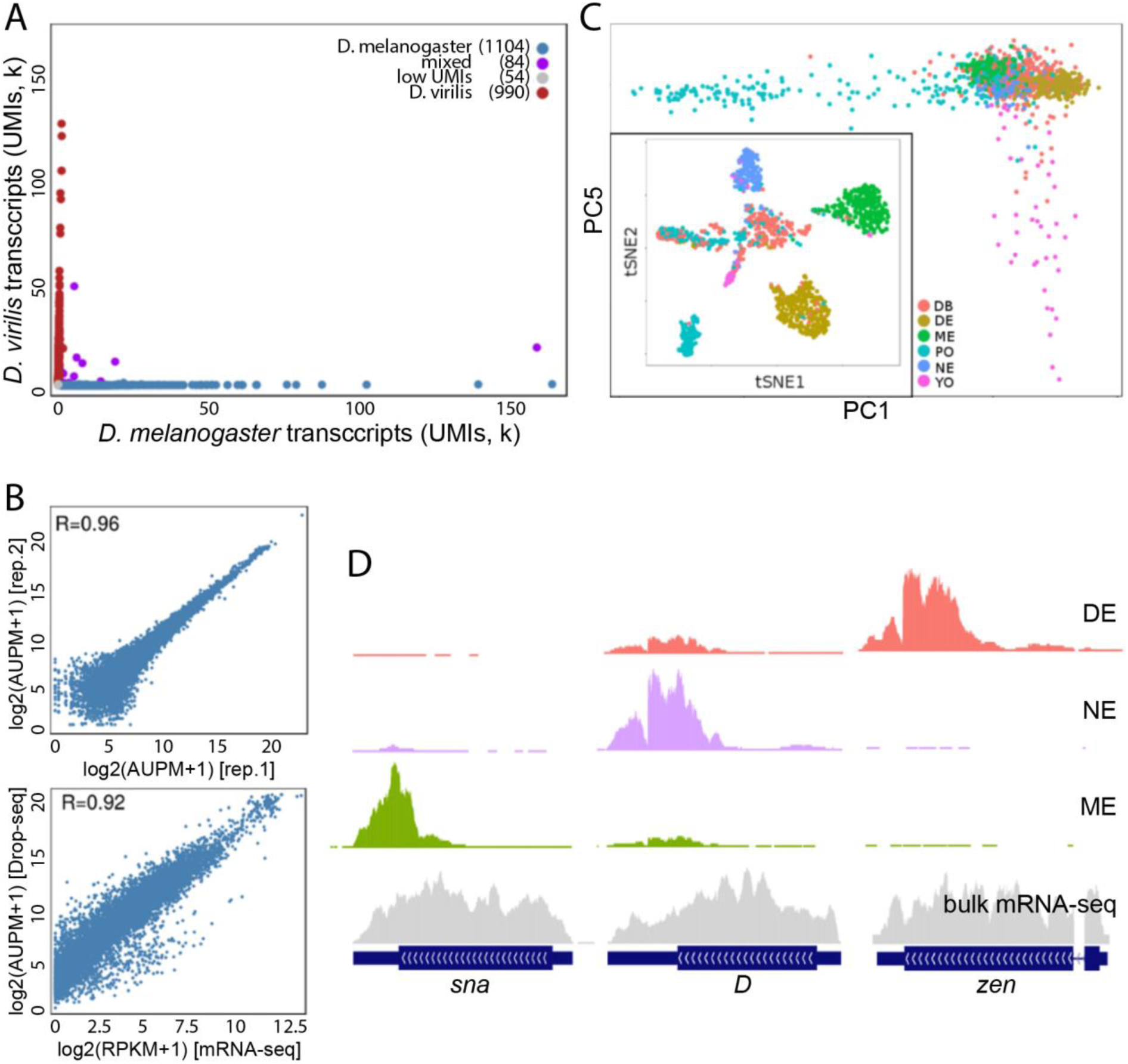
Drop-seq reproducibly resolves single-cell transcriptomes and identifies distinct cell populations. (A) Species separation plot for one of the *D.mel–D.vir* replicate. The mixed species doublet ratio is < 4%. Same species doublets would be expected at a similar rate. Grey colored cells contain less than 1 000 UMIs. (B) Correlation of aggregate gene expression between independent Drop-seq replicates (top) and between Drop-seq replicates and bulk mRNA sequencing from live and intact embryos; Pearson correlation shown on the top. (C) Outset: principal component analysis of 1 369 cells showing separation of pole cells (PO) and yolk cells (YO) along PCs 1 and 5. Cell types colored as indicated according to marker gene expression, or being identified as a doublet (DB). Inset: t-SNE analysis of the same cells shows clustering of cell types as indicated. Doublets expressing multiple marker classes cluster centrally. (D) Genome browser tracks at 3 genes, expressed in DV domains. Whole embryo mRNA sequencing is shown (bulk, grey), as are the aggregate transcriptomes of cells expressing dorsal ectodermal (DE, red), neurectodermal (NE, purple), or mesodermal (ME, green) marker genes (*in silico* tissue dissection).

For all replicates, we observed that the cumulative fraction of reads as a function of the number of sequenced cells had a well-defined inclination point (‘knee’, see Suppl. Fig. S1). We defined cutoffs for the number of UMIs per cell. On average, each run yielded ~1 546 cells containing an average median of ~2 040 genes/cell and ~5 430 UMIs/cell (excluding mixed-species events, Suppl. Table S3). By essentially reconstituting the embryo *in silico* (merged UMIs), we found that Drop-seq replicate correlation was generally high at R > 0.94 (Fig 2B, Suppl. Fig S1B), as was correlation between Drop-seq samples and stage-matched whole unfixed embryos that were poly-A sequenced in bulk with R > 0.88 (Fig 2B, Suppl. Fig S1B). Comparison of the expression levels of individual genes (UMIs merged across stamps) and absolute quantification of selected genes in individual stage-matched embryos by N-Counter *(33)* shows that the positive quantitative correlation holds true for tissue-specifically expressed genes. (Suppl. Fig. 1C). We conclude that Drop-seq data accurately reflects the transcriptomic state at the embryo level, which indicates that individual cells are sequenced at random without bias for any particular cellular identity. Furthermore, principle component analysis shows that cells do not separate along principles components by biological replicate (Suppl. Fig S1F), which underscores the technical reproducibility of Drop-seq.

### *In silico* dissection of the embryo

Principal component analysis was performed on the transcriptomes of *D. mel.* cells that scored high regarding DE, NE and ME markers (Sup. Fig. 1E), and were identified as doublets (Sup. Fig. 1D), pole, or yolk cells (Sup. Table. 4). The analysis reveals several very distinct cell populations (Fig. 2C). For example, the first principal component clearly separates pole cells (Fig. 2C, blue) according to abundant and distinctive expression of specific marker genes such as *primordial germ cells* (*pgc*) (Suppl. Table 4). Pole cells constitute the *Drosophila* germ line and are the first to cellularize at the posterior dorsal tip of the embryo at stage 5. They constitute a discrete cellular lineage from stage 5 onwards and contribute to no other developmental fate (*34*). Similarly, yolk nuclei primarily play a supportive role aiding in yolk digestion and energy metabolism, but do not participate in patterning (*35*). Marker analysis shows that yolk nuclei separate along PC5 (Fig. 2C, pink). t-distributed stochastic neighbor embedding (t-SNE), which reduces the dimensionality of the 7 975 single cell transcriptomes along principal components lends further support for the distinctiveness of the pole cells and yolk nuclei, as both form coherent clusters (Fig. 2C inset). Other clusters that are readily apparent are cells specifically expressing DE marker genes (Fig. 2C inset, tan cluster), ME marker genes (inset, green cluster), or NE marker genes (Fig. 2C inset, pale blue cluster). Hence, both PCA and t-SNE readily separate cells according to biologically meaningful patterns, including positional DV identity.

An immediate benefit of the single cell sequencing data is that it enables *‘in silico* dissection’ of the embryo – based on marker gene sets, individual cells can be identified and their average gene expression profiles can be generated by merging sequencing reads from defined sets. We first examined the exclusivity of marker genes that are expressed in largely abutting domains along the DV axis of the embryo (Suppl. Table 4) on a cell-by-cell basis. Ternary analysis shows that cells expressing ME, NE, or DE marker genes are well-segregated from each other (Suppl. Fig. S1E), which highlights the specificity of the marker gene sets and the faithfulness of the single cell sequencing data. We aggregated the single cell transcriptomes of these 3 marker-based sets of cells into regional expression profiles. The resulting transcriptome tracks accurately recapitulate the exclusivity of known marker genes; for example, *snail, Dichaete* and *zerknüllt* are specifically expressed in the ME, NE, or DE, respectively (Fig. 2D). In effect, we conducted a marker-based dissection of the intact wild type embryo and obtained tissue specific transcriptomes – something not possible with conventional methods. A major benefit is that the cell population of interest could be refined by any combination of marker genes and even by thresholding on expression levels, for example.

For further analysis, we focus on the cells most likely to harbor unambiguous patterning information. As such, we eliminated pole cells as well as yolk cells from further consideration because they constitute distinct lineages that do not shape embryonic patterning of the embryo. Additionally, we aimed to identify potential doublets to avoid inconsistent transcriptome information. Cells within the central t-SNE cluster (Fig. 2C inset, salmon) express several marker genes belonging to different DV domains (Suppl. Fig. S1D) and likely represent same-species doublets (~2%). Furthermore, we only considered cells with ≥12 500 UMIs to assure sufficient patterning information (see Materials and Methods for details). The remaining highest-confidence set encompasses ~1 300 cells (‘high quality cells’, HQCs) with a median unique transcript number of > 20 800 UMIs/cell mapping to 3 131 genes/cell (Fig 3A). It should be noted that the sequencing depth per cell is substantial for single cell approaches, in particular given that the *Drosophila* genome is comparatively small with only ~14 000 coding genes. Principal component analysis of the HQC transcriptomes (Fig. 3B) readily identifies the biological variation due to germ layer (DV) identity along the first two principal components, with ME, NE, and DE marker genes clearly separating. Additionally, we scored the cells expressing DsRed, a transgene present in the sequenced embryos under control of a characterized *vnd* enhancer (36) that directs expression in the ventral neurogenic ectoderm (Suppl. Figure S2A). DsRed transcripts are detected in a subset of cells that also score highly for NE markers and sequester in an NE subcompartment in principle component analysis (Fig. 3B), further underscoring that Drop-seq data faithfully resolves cell-specific transcriptomes.

**Figure 3.**
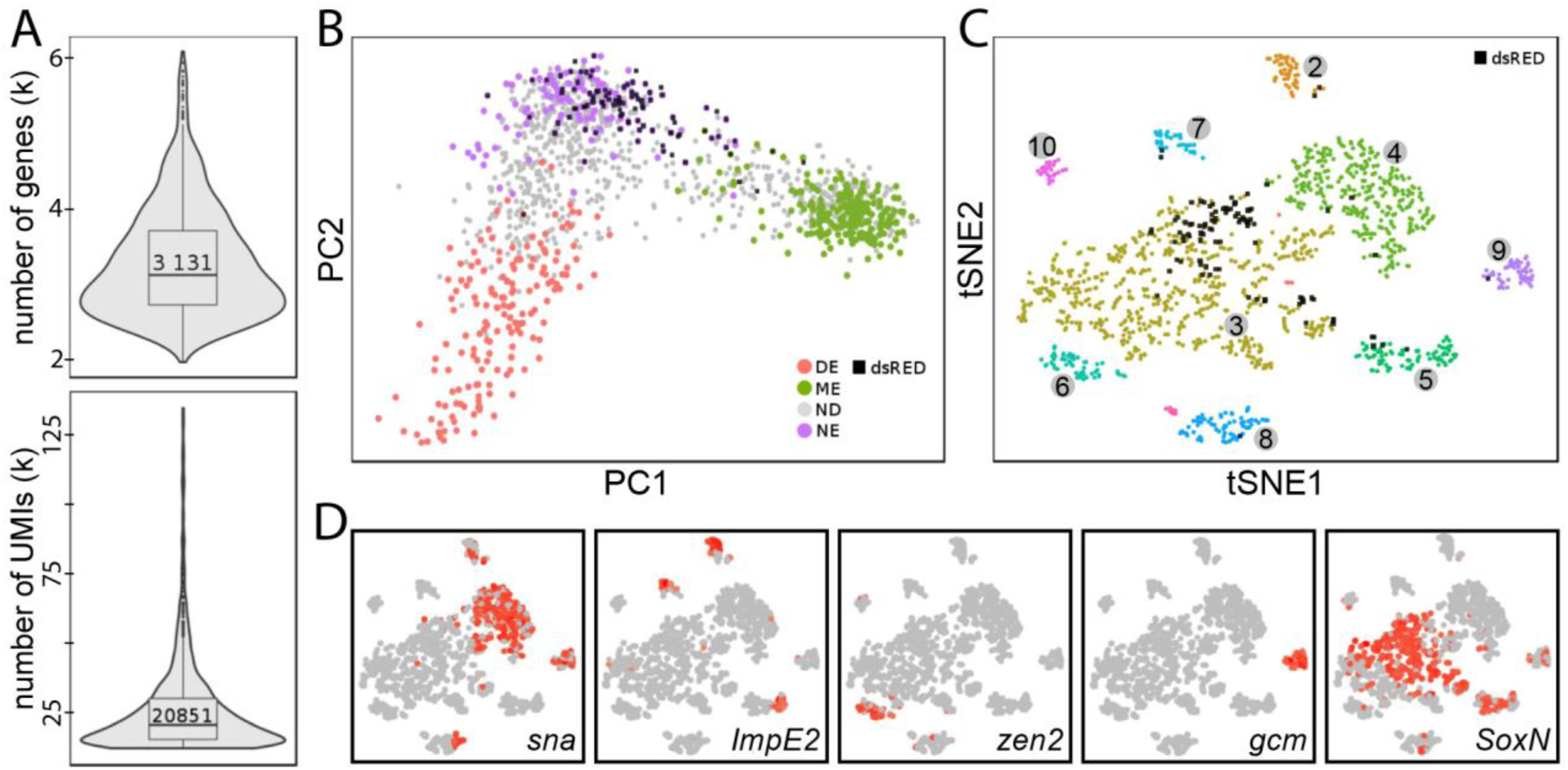
High-quality transcriptomes resolve the embryo spatially. (A) Violin plots showing the distributions of gene (top) and UMI (bottom) number detected per cell for the ~1 300 highest quality cells used for spatial mapping. (B) Principal components PC1 and PC2 separate DE, NE, and ME. Color assigned according to marker genes; grey cells could not be assigned. Black boxes indicate cells highly expressing the *vnd::DsRed* reporter gene and are correctly located in the NE population. (C) t-SNE analysis clusters the ~1 300 cells based on highly variable genes into major clusters. (D) Examples of highly variable genes specifically expressed in t-SNE clusters. These gnenes are known to have specific patterned expression, indicating that t-SNE clusters resolve spatially. Cells expressing the indicated genes highly are marked in red.

### Single cell clustering resolves spatial identities

t-SNE analysis of the HQCs identifies at least 10 coherent clusters of cells isolated from stage 5/6 embryos (Fig. 3C). Among the most specific genes that drive t-SNE clustering are many that are well-known for their roles in embryonic patterning and tissue specification (Suppl. Table S5). For example, the gene *sna* encodes a Zn-finger transcription factor (TF) known for its early requirement in ventral mesoderm development as it represses more dorsal fates; *sna* is among the 15 genes most specific for cluster 4. Similarly, the homeodomain TF *zen2* is known to be expressed dorsally in regions that give rise to the dorsal epidermis and extra-embryonic membranes, whereas the SOX TF *SoxN* is expressed broadly and laterally in the presumptive neurogenic ectoderm; the genes *SoxN* and *zen2* are among the most specific genes for clusters 3 and 6, respectively. In fact, the most specific genes (driver genes) within each cluster are specifically expressed in overlapping spatial domains, and while some of those cluster-specific expression domains are patterned along the DV axis, others are patterned along AP. Thus, t-SNE clusters, which group cells according to transcriptome similarity, appear to reveal basic patterning domains within the embryo. However, the t-SNE cluster calling shown in Fig. 3C demonstrates that it does not fully resolve the spatial organization of the embryo – even within clusters such as cluster 3, finer organization is apparent. For example, when scoring cells for expression of the *DsRed* reporter gene (Fig. 3C, black squares), *DsRed* is clearly enriched in a subset of cluster 3, which is spatially confined in the t-SNE map. This is in agreement with genes like *SoxN* marking the entirety of the NE, while the *vnd* enhancer drives *DsRed* expression only in the ventral-most third of the NE. While clusters are defined by driver genes, their expression is not cluster-exclusive: for example, *SoxN, sna,* and *zen2* are drivers of clusters 3, 4, and 6, respectively, their expression can also be detected in cells of other clusters (Fig. 3D). This demonstrates that clustering reveals transcriptome similarity according to multiple patterning cues (e.g. along the AP and DV axes), thus resolving a more finely grained spatial identity of participating cells.

Four observations regarding the most highly cluster-specific genes (Suppl. Table S5) should be noted: (i) A majority of cluster drivers encode TFs (which is not unexpected). (ii) Several of the highly variable genes remain un- or understudied in early development (e.g. *DNaseII, Z600, Meltrin, mtt, Atx-1,* as several CGs), though their specific expression might implicate them in fundamentally contributing to cell identity. (iii) Eight of the most variable genes (*fkh, kni, grn, hb, Abd-B, ken oc,* and *toy,* see Suppl. Table S5) are TFs that occur among the most variable genes in > 1 cluster, highlighting the impact and importance of combinatorial gene regulation for cluster and cell identity; and (iv) a surprising number of genes among the most variable genes are noncoding RNAs (*CR45361, CR44683, CR43302, CR43279, CR41257, CR45185, CR45185*), which may indicate a more central role for lncRNAs in early embryonic patterning and development.

When considering GO-term enrichment among genes up-regulated in specific clusters, all are enriched for terms indicating transcriptional regulation (Suppl. Fig. S3A). In fact, the only non-transcriptional regulation related terms are growth factor signaling terms in cluster 3. This supports the spatial association of cluster 3 in lateral domains, as the NE is a source of FGF and EGF signaling molecules (*37, 38*). Considerably more complex and broadly informative are GO-term enrichments for biological function (Suppl. Figs. S3B, S4). For example, cluster 3 was enriched in terms relating to the nervous system (*ventral cord, neuroblasts, neurogenesis, CNS*). Similarly, mesodermal terms are primarily associated with cluster 4, placing those cells ventrally. Terms referring to the extra-embryonic membranes (*amnioserosa, dorsal closure*) should place cells in cluster 6 dorsally, while hindgut terms should place cluster 8 cells in posterior regions. Though these terms generally highlight the clusters’ likely spatial association within the embryos, enriched terms often do not lend themselves to simple spatial assignment – cells belonging to a single cluster may be associated with terms indicative of several distinct regions, and separate clusters can be associated with terms indicative of the same region. Clusters 3, 6, and 8, for example, are all enriched for “*generation of neurons*”, which are likely lateral terms; however, cluster 6 is also enriched for genes associated with "heart development” (likely mesodermal) and “amnioserosa” (dorsal ectoderm), while cluster 8 is enriched for genes annotated to be involved in *“head development*” (anterior) and “*hindgut*” (posterior) (Suppl. Fig S3B). Naturally, attempting such spatial inferences pushes the information value of GO terms beyond their intended limits, but it does demonstrate that spatial information is captured by clustering. To truly understand the spatial relationship of transcriptional programs, it was necessary to map cells back onto the embryo, and we aimed to do that solely based on the single cell transcriptomes.

### Reconstructing the embryo by computational spatial mapping

Motivated by previous efforts in the early zebrafish embryo and the central nervous system of the marine annelid *Platynereis dumerilii* (*18, 19*), we reasoned that given our single cell transcriptome data and a sufficient number of gene expression patterns resolved at the single cell level, we should be able to map cells back into the embryo; the degree of mapping confidence should be directly dependent on the accuracy and the combinatorial complexity (entropy) of the reference mapping dataset. If successful, a virtual embryo would emerge where all spatial positions can be queried for gene expression (Fig. 1B). The Berkeley *Drosophila* Transcription Network Project (BDTNP) has registered *in situ* hybridization data for the relative spatial expression of 84 individual genes in the early embryo, resulting in a quantitative gene expression reference map of a virtual embryo (*28*). We wondered whether given the spatially and quantitatively resolved expression profiles of 84 marker genes measured across 6 000 cells (positional bins) by the BDTNP, we would be able to accurately place randomly selected positional bins back at their original location. To minimize the combinatorial complexity, we first binarized gene expression into ON and OFF states for each of the 84 genes and each of the 6 000 positional bins in the BDTNP reference atlas (Fig. 4A, I); the binarization thresholds were chosen manually and for each gene independently to recapitulate *de facto* expression as revealed by RNA *in situ* hybridization (*25, 26*). We find that the positional information encoded among the 84 mapped genes is sufficient to map single-cell transcriptomes back to their original location within 2-3 cell diameters (Fig. 4B, green). When considering all 3 000 bilaterally symmetric positional bins, we find that our mapping coverage is very high throughout the embryo (Fig. 4C, blue regions), with few small regional exceptions where the embryo coverage is below average (Fig. 4C, white regions).

**Figure 4:**
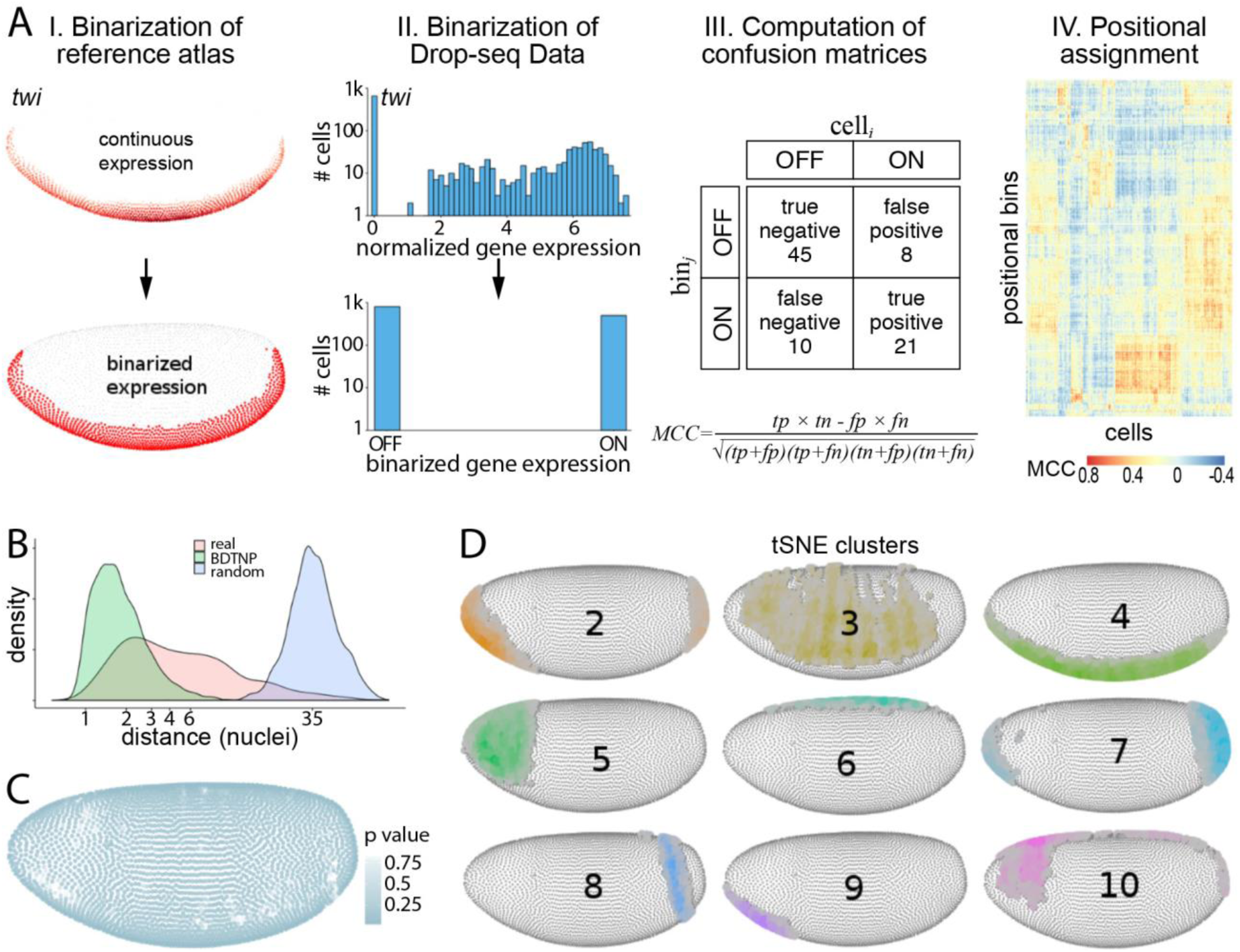
Reconstruction of the *D. melanogaster* embryo. (A) Mapping procedure overview. (I) The 84 digitized BDTNP gene expression patterns were binarized. (II) Gene expression across the ~1 300 cells was binarized to maximize marker gene correlation guided by the BDTNP reference atlas. (III) Confusion matrices are calculated scoring expression (dis)agreement between the binarized transcriptomes and the ~3 000 positional bins of the reference atlas. The Mathews correlation coefficient (MCC) is calculated for every cell and every bin. (IV) Positional assignment for each cell is diffused, based on its MCCs across all bins. (B) Density plot showing mapping confidence in terms of mean Euclidean distance between a cell's highest scoring location and the following six ones. Drop-seq data shown in red; mapping of BDTNP bins onto themselves (green) and mean Euclidean distance between random bins (blue) serve as positive and negative control respectively. (C) Mapping coverage across the embryo. More than 87% locations of the embryo are covered (blue regions), as their highest score is significantly higher than those obtained by chance (p-value < 0.05, see Suppl. Materials and Methods for details). (D) Averages of diffused scores of cells within t-SNE clusters 2-10 reveal spatially localized cell populations.

To map each of our 1 300 high quality transcriptomes to positional bins, we binarized the Drop-seq data to the BDTNP reference atlas (Fig 4A, II) (see Suppl. Materials and Methods for details). We then compared the expressions of each cell across all positional bins and collected the (mis)matches into confusion matrices, which allowed us to assign cell-bin scores by computing the Matthews correlation coefficients (MCCs) (Fig 4A, III). The result is a diffused mapping score for any given sequenced cell across all positions in the embryo (Fig. 4A, IV). Fig. 4B demonstrates that sequenced cells are mapped to a territory of only a few cell diameters with high confidence (Fig. 4B, red), whereas mapping positions for cells with randomized scores are spread throughout the whole embryo (Fig. 4B, blue). Our mapping algorithm allows us to compute the spatial expression of a gene at any position by carefully combining the normalized gene expression with the MCC scores (see Suppl. Materials and Methods for details). Doing this separately for all positional bins produces what we call a “virtual *in situ* hybridization” (vISH) (e.g. Fig. 5A).

**Figure 5:**
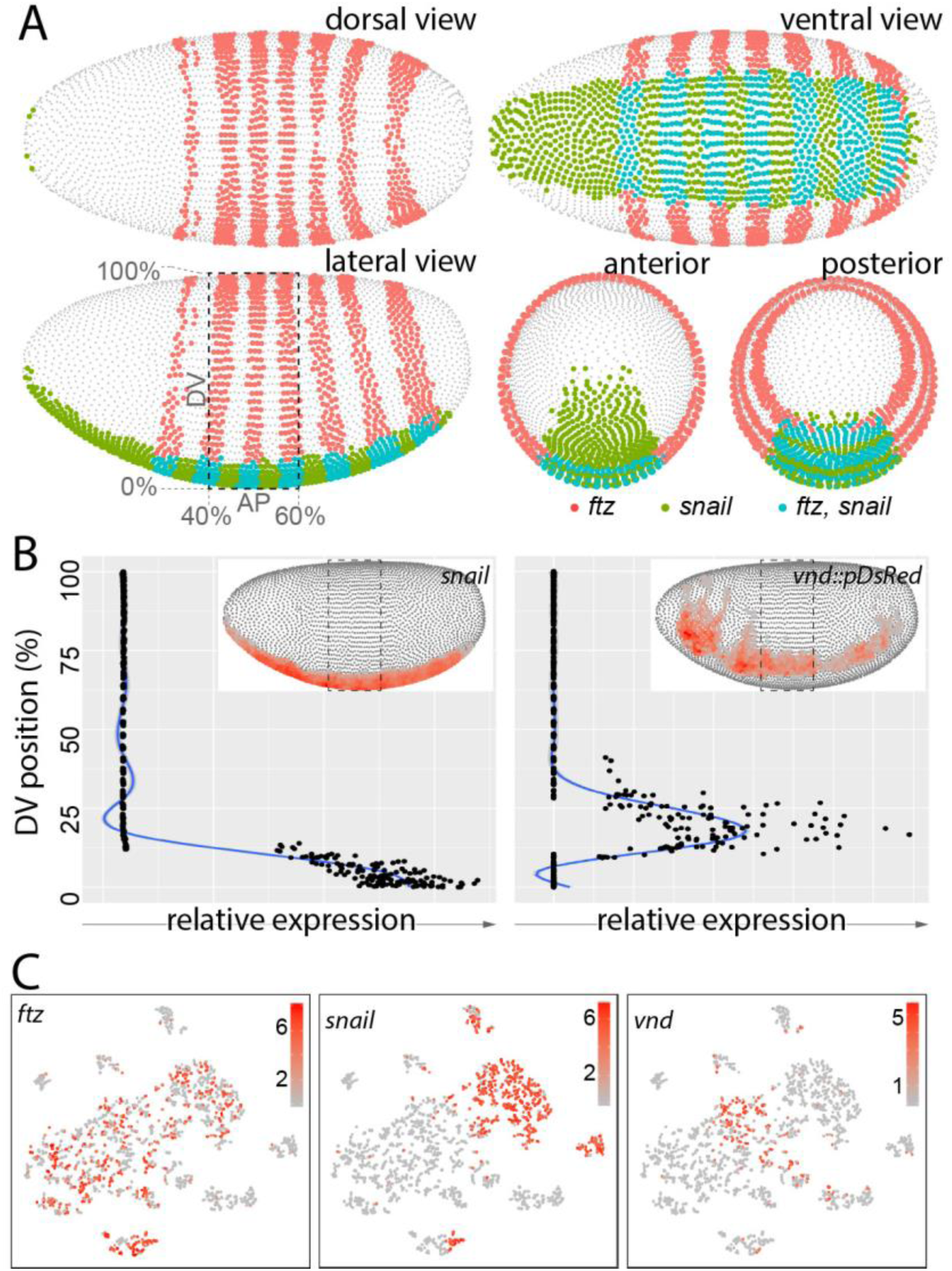
*Drosophila* Virtual Expression eXplorer (DVEX). DVEX (www.dvex.org) is an online resource allowing visualization of mapped single cell sequencing data. (A) Virtual in situ hybridization (vISH) for 2 sample genes, the pair rule gene *ftz* and the mesodermal gene *sna* (red and green, respectively, overlap in cyan) in five representative views. (B) Expression along the DV axis of *sna* (left) in the ventral mesoderm and the *DsRed* transgene under control of a *vnd* enhancer (right) in the ventral NE; the *vnd* enhancer is known to be repressed by *sna.* DVEX allows for similar visualization along the AP axis. (C) Cells expressing individual genes (*ftz*, *sna*, *vnd*) marked in t-SNA cluster context; expression level (relative UMIs) shown in red.

Calculating the average mapping scores per bin across all cells belonging to a cluster produces a spatial signature for each cluster (Fig. 4D). Accordingly, cells of clusters 2, 8, and 7 map along the AP axis to terminal and distinct sub-terminal regions, while clusters, such as 3, 4, and 6 map along the DV axis to ventral, lateral and dorsal regions, respectively. Several clusters appear to be the result of converging AP and DV patterning mechanisms: clusters 5 and 9 are both found in anterior regions, but while 9 is exclusively ventral, 5 appears only lateral and dorsal. Interestingly, cluster identity and mapping position is not mutually exclusive. For example, cells of clusters 4 and 9 map to overlapping ventral domains, which may suggest substantial regional transcriptomic heterogeneity not previously appreciated.

### vISH accurately predicts spatial gene expression

We have mapped each of the 1 300 HQCs to the embryo, allowing to resolve the local transcriptome state. The ***D**rosophila* **V**irtual **E**xpression e**X**plorer (DVEX) tool provided online (www.dvex.org), allows the generation of vISHs for any gene detected. vISH patterns for single or even multiple genes can be queried and displayed on a virtual embryo in multiple views (dorsal, lateral, ventral, anterior and posterior, see Fig. 5A); a threshold selection option allows the user to adjust the mapping scores to be visualized. Additionally, expression gradients can be estimated (Fig. 5B) along the AP and DV axes. Furthermore, DVEX provides an interactive environment to explore the t-SNE-generated clusters, genes that drive the clustering, and which cells in which clusters express any gene of interest (Fig. 5C).

We generally observe close concordance between vISH predictions and de facto expression as detected by RNA *in situ* hybridization. This holds true for marker genes expressed in a wide variety of AP and DV patterns, such as *stumps* expression in the mesoderm, *SoxN* in the lateral ectoderm, *gcm* in an anterolateral patch, *Oaz* in an antero-dorsal patch, *Kruppel* in a gap pattern and *eve* in a primary pair rule pattern (Fig. 6A, Suppl. Fig. S5). For several genes where ISH images were not available, we have made *de novo* predictions by vISH and confirmed their accuracy by RNA ISH. For example, our *dsRED* transgene was correctly mapped to the ventral neurogenic ectoderm (Fig. 6B and Suppl. Fig S6, also see Suppl. Fig. S2). *CG4500* appeared strongly expressed by virtual embryo dissection in cells positive for mesodermal markers and shows a vISH pattern in the presumptive mesoderm; this is in concordance with RNA ISH (Fig. 6B, S6). Similarly, vISH predictions were in agreement with measured *in vivo* expression for the weakly expressed genes *ana* in the dorsal ectoderm and *babos* in a stripe-modulated pattern in ventral regions with stronger expression anteriorly than in the trunk of the embryo (Fig. 6B, S6). We were especially surprised by the vISH accuracy for *CG16886,* which was projected to be expressed in 3 small dorsal patches. In fact, we detected specific and strong expression as predicted (Fig. 6B, S6), highlighting the signal:noise advantage of single cell sequencing, as signatures are not diluted over populations of cells. Furthermore, *CR45693,* a non-coding lncRNA was detected in our single cell data. We noted that it appeared to be expressed only weakly; in fact, it was not called as expressed in early whole embryo sequencing data (*29*). However, vISH as well in silico dissection predicted it to be expressed in dorsal regions (Fig. 6B, S6). Indeed, RNA ISH against *CR45693* shows weak, but detectable and reliable expression with specific cytoplasmic signal in a dorsal stripe at stage 5/6 (Fig. 6B, Suppl. Fig. S6). Similarly, we could confirm a second lncRNA, *CR44691,* to be weakly expressed in the dorsal ectoderm as predicted (Suppl. Fig. S5). A third lncRNA expression prediction was only partially confirmed. The lncRNA *CR44917* was expected by vISH to be stripe modulated with expression in 2 AP stripes in lateral and dorsal regions, as well as a posterior patch (Suppl. Fig. S6). The RNA ISH showed weak general expression, but we could neither confirm the anterior stripe nor the posterior patch expression. Only the post-cephalic stripe and the mesodermal exclusion aspects of the predicted pattern appear to hold true (Suppl. Fig. S6). A further discrepancy between detected (i.e BDTNP measured) and predicted expression we observed was for the gene *rhomboid* (*rho*). The measured expression was present in 3 domains, 2 lateral stripes in the ventral NE and a single stripe in the DE (Suppl. Fig. S2B, top). Predicted *rho* expression by vISH showed the same general 3 domains, but the lateral stripes were noticeably more ‘patchy’ and modulated along AP (Suppl. Fig. S2C, right). We assessed *rho* expression by RNA ISH more carefully and found that *rho* expression is highly dynamic at stage 5-6 – early stage 5 exhibits near uniform lateral stripes along AP, but this pattern quickly becomes stripy and patchy by stage 6 (Suppl. Fig. S2C). Thus, the vISH prediction was not only in concordance with observed *rho* expression, but the fact that a more uniform *rho* pattern was part of the BDTNP mapping atlas indicates that the mapping algorithm using a set of 84 measured expression guides does not force pattern predictions to conform to mapping guides. Taken together, we demonstrate that single cell transcriptome data once mapped back to its point of origin can reliably predict gene expression patterns and aid in the identification of new marker genes with distinct spatial behaviors. This should prove especially useful where expression is weak or spatially limited. As we can readily produce a vISH for every gene detected in our data, we predicted the spatial expressions of the 476 most highly varied genes. We clustered their correlation matrix and identified 10 parental branches (Sup. Fig. 7). Averaging the vISHs of the genes within each branch generated archetypes of expression patterns, spanning the different spatial domains (Sup. Fig. 7), which enables a comprehensive classification of genes falling within an archetypal expression.

**Figure 6:**
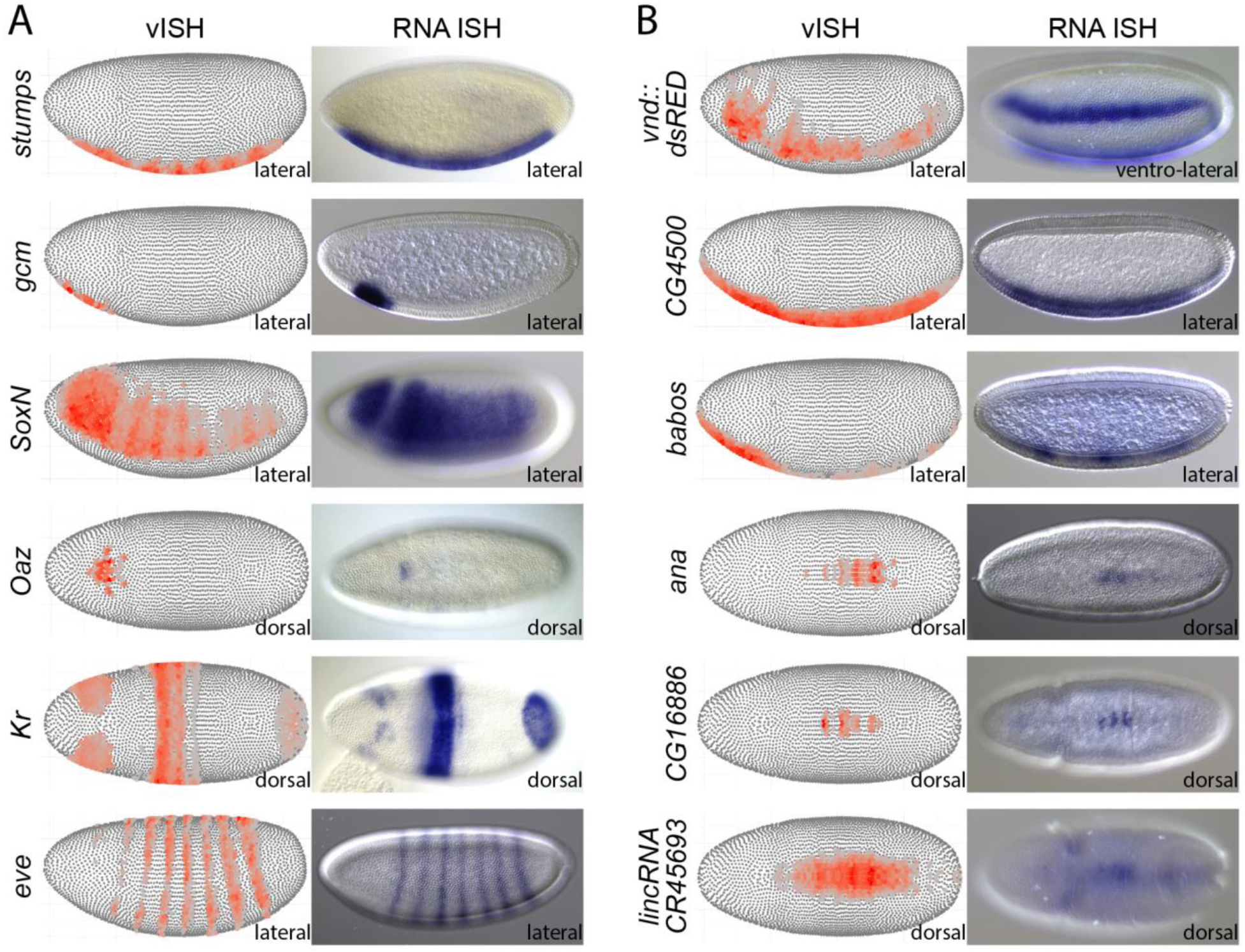
vISH accurately predicts spatial gene expression patterns. (A) vISH predictions of genes with known expression patterns (left) and *in vivo* expression by in situ hybridization (right). Embryo orientations anterior left, DV rotation is indicated. (*Kr, stumps, Oaz* images from BDGP). (B) *De novo* spatial expression predictions by vISH (left) and *in vivo* validation by *in situ* hybridization (right). Embryo orientations anterior left, DV rotation is indicated.

**Figure 7:**
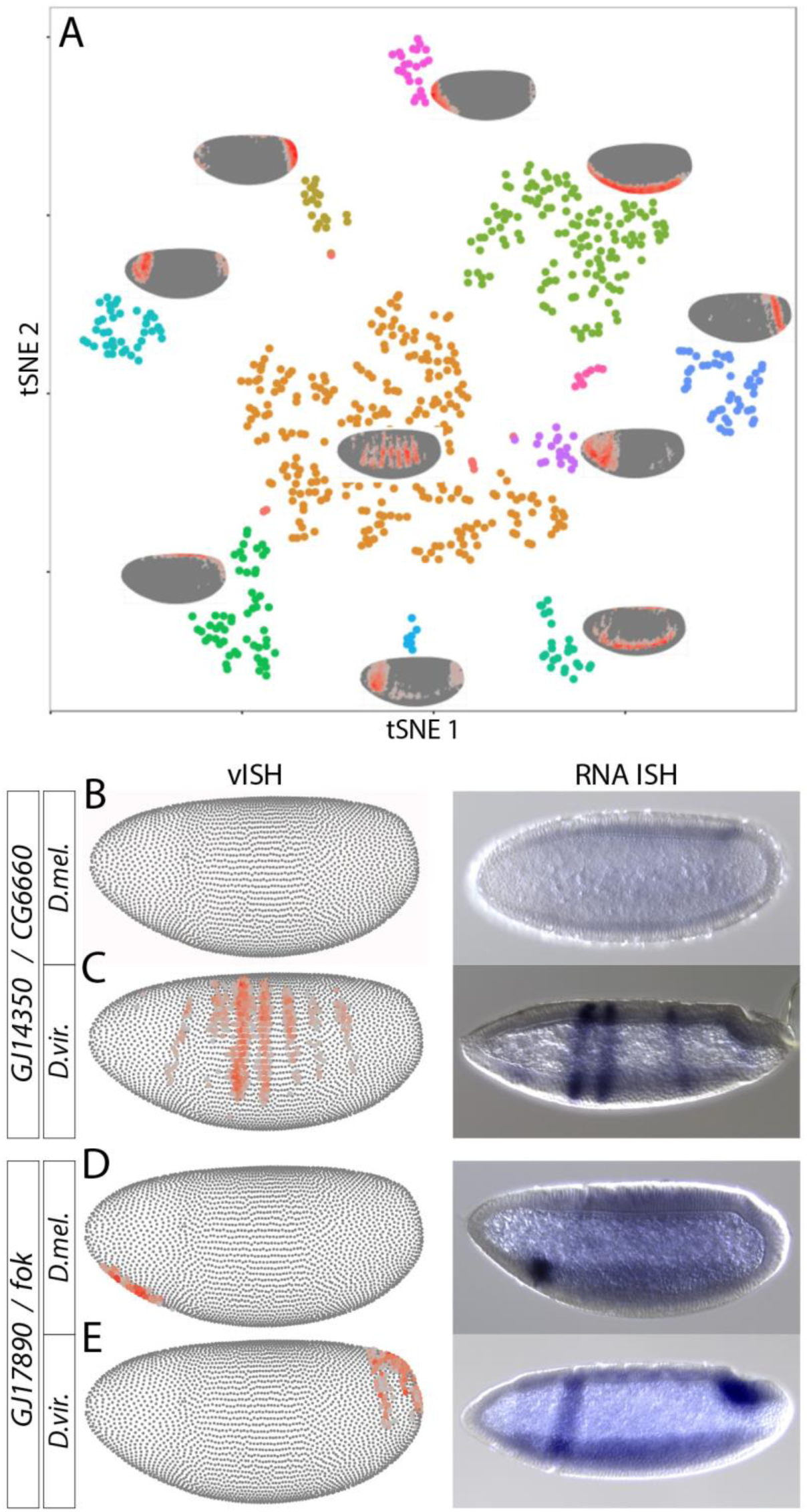
vISH detects evolutionary changes in gene expression patterns. (A) t-SNE clustering of cells from stage 6 *D. vir.* embryos; spatial localization of each cell population on the embryo is indicated as in Fig. 4D. (B - E) Gene expression divergence between homologous gene pairs predicted by vISH (left) and experimentally validated by RNA ISH (right) for the *D. mel.* genes *CG6660* (B) and *fok* (D) and their respective *D. vir.* homologs *GJ14350* (C) and *GJ17890* (E).

### Evolutionary changes in gene expression pattern can be detected by vISH

2 847 *D.vir.* cells were sequenced at a depth of ~4 780 unique transcripts/cell and ~1 217 genes/cell. After eliminating yolk, pole cells, and doublets, we were left with a much sparser dataset of 673 HQCs, which we clustered and mapped as before (Fig. 7A). The clustering generated for *D.vir.* (Fig. 7A) bears a striking similarity to that of *D. mel.* with respect to cluster number, proportional cluster sizes, as well as where these clusters map onto a virtual *Drosophila* embryo (compare Fig. 2E, 3C with 5B). Correlation of gene expression between merged transcriptome data of both *Drosophilids* was high (Pearson R = 0.77). While the vast majority of vISHs generated for the respective homologous genes agree in terms of expression prediction, we searched for genes that change vISH-predicted expression between the two species. A small number of genes was found to be expressed in one, but not the other species, while another small number of genes were predicted to change expression patterns between species. Though these vISH comparisons are based on relatively few sequenced cells, we tested the putative expression divergence of the gene pairs *CG6660 / GJ14350* and *fok / GJ17890.*

*GJ14350* and *CG6660* were determined to be homologous based on protein conservation and synteny (Suppl. Table S7), but while *CG6660* was predicted to not be expressed in *D. mel.* (no UMIs detected) (Fig. 7B, left), its homolog *GJ14350* was reasonably abundant (576 UMIs total) in *D.vir.* single cell sequencing data. vISH predicted *GJ14350* to be expressed in an AP stripe modulated pattern, primarily in two stripes posterior to the cephalic furrow (Fig. 7C, left). We found no detectable expression for *CG6660* at stage 6 (Fig. 7B, right), whereas its homolog *GJ14350* was expressed in several stripes and most strongly in two stripes (Fig. 7C, right), as predicted. For the homologous pair *fok / GJ17890* (see Suppl. Table S7), *fok* was predicted and verified to be expressed in an anterior ventral patch in *D.mel.* (Fig. 7D), whereas *D. vir.* vISH suggested absence of the anterior patch and weak expression posteriorly. While there is a tendency for low posterior expression as early as stage 6/7, this is highly variable and robust posterior staining is not seen until stage 8. However, the absence of the specific anterior staining in *D. vir.* was clearly confirmed (Fig. 7E). Therefore, vISH can serve as an invaluable resource for the prediction of evolutionary changes in spatial gene expression dynamics.

## DISCUSSION

Here we have disassembled the stage 6 *Drosophila* embryo, sequenced its constituent cells, and reconstructed the embryo to arrive at the transcriptomic makeup of a ~6 000 cell tissue with unprecedented spatial resolution. Essentially, the virtual embryo that emerges is a gene expression blueprint of embryonic development at unprecedented resolution – single cell transcriptomes mapped onto a 3D lattice, which can be queried in context: *Which genes are co-expressed? Which are exclusive? Which cells invoke signaling cascades? How do surrounding cells respond?.* We show that the expression data faithfully recapitulates know patterning domains, qualitatively and quantitatively. Single cell transcriptome data generally allows for *in silico* dissection of specific cell types based on marker genes, such as the average transcriptomes of even (eve+) vs. odd (odd+) pair rule stripes, for example – a dissection that would not be possible physically *or* genetically. An *in silico* dissection along the DV axis conforms extremely well with known spatial gene expression limits, but further filtering by markers combinatorially would allow for an ever finer-grained analysis only limited by the number of cells sequenced. Cell types that can not develop properly without the correct tissue context, are thus finally accessible for genomic studies via single cell sequencing. From a gene regulatory perspective, single cell transcriptomes make it possible to accurately distill rules of gene expression – gene sets that are, for example, concomitantly or mutually exclusively expressed can be defined, which opens a new avenue to extract regulatory rules in complex tissues.

Using known markers, we were able to confidently map single cells to their most probable embryonic positions of origin, which in turn allows for the accurate prediction of previously untested gene expression patterns. Due to the improved signal: noise ratio compared to whole embryos, expression that is characteristic to only a small subset of cells can be reliably detected and projected onto the regions of origin. The *Drosophila* germ line (pole cells) for example proved to be a small but highly distinct cell population, which would be severely diluted in whole embryo studies; in fact, pole cells were so fundamentally different in terms of their transcriptome, that we decided to exclude them from further analysis early on, simply to avoid them skewing any comparative approaches. The capability of rigorously identifying features shared by only a small subset of cells may be especially important, because several of the lncRNAs (which are finally accessible) show not only highly specific expression in, for example, distinct t-SNE clusters, but are often expressed rather lowly. Given recent studies regarding their roles in gene and genome regulation from the regulation of TFs to genome organization, reliably identifying and characterizing lncRNAs will be crucial. The approach used here combines the unbiased nature of whole genome sequencing with the spatial resolution of *in situ* hybridization, which opens the transcriptional output of the coding as well as the non-coding genome to systematic interrogation.

As individual cells, rather than pools, become the focal points of investigation, entirely new questions about how a tissue becomes organized are now possible, starting with understanding tissue heterogeneity, but leading to questions about how cells communicate and behave in spatial relation to each other. We show that comparative approaches can distill evolutionary changes in spatial gene expression from mapped single cell transcriptome data, but the same analysis will enable unprecedented insights into the cell-specific responses to environmental, chemical, and genetic challenges.

## AUTHOR CONTRIBUTION

NR had the initial idea; NR, RZ defined the broader strategy and supervised; NK, PW, JA, CKo, NR, RZ designed the experimental and analytical strategy; PW did the fly genetics and embryo collections; PW, CKi, CKo, prepared the biological materials; PW, JA, AB, SA, CKo, performed Drop-seq and prepared sequencing libraries; NK, PW, CKo, NR, RZ analyzed data; NK performed almost the entire computational analyses, in particular designed and implemented mapping algorithms, their parameterization, optimization and visualization; PW, CKi validated predictions experimentally; NK designed and implemented the interactive DVEX browser interface; NR, RZ procured funding; NK, PW, CKo, NR, RZ wrote the manuscript.

## ACKNOWLEDGEMENTS

We are indebted to Mark Biggin (LBNL) for fruitful discussions, technical advice and sharing of unpublished BDTNP data, Soile Keranen (LBNL) and Sarah Ugowski (MDC) for experimental assistance. We would like to thank Steve Small (NYU) for sharing transgenic *Drosophila* lines, Angelike Stathopoulos (Caltech, NIH R35GM118146) for sharing unpublished results and discussions. Nir Friedman (Hebrew University), Eileen Furlong (EMBL), David Garfield (HU), Jan-Philipp Junker (BIMSB/MDC), and John Rinn (Harvard), as well as members of the Rajewsky and Zinzen labs for constructive discussions. Dan Munteanu (MDC) for IT support. The Max Delbrück Centre and the DFG (SPP 1738, RA 838/8-1, RA 838/5-1) for funding.

## MATERIALS AND METHODS

### Fly Strains and *in situ* hybridizations

The *Drosophila* strains used were *D. virilis w^e^* (*Drosophila* Species Stock Centre, # 15010-1051.17) *and D. melanogaster y^1^ w^1118^; P{st.2::Gal4}; P{vnd::dsRED},* where *st.2::Gal4* was crossed in from a kind gift by Steve Small (NYU) and *vnd::dsRED* was created by standard P-element transgenesis (*1*) after placing the early *vnd* enhancer (*2*) into *pRED-HStinger* (*3*).

**Suppl. Table S1:**
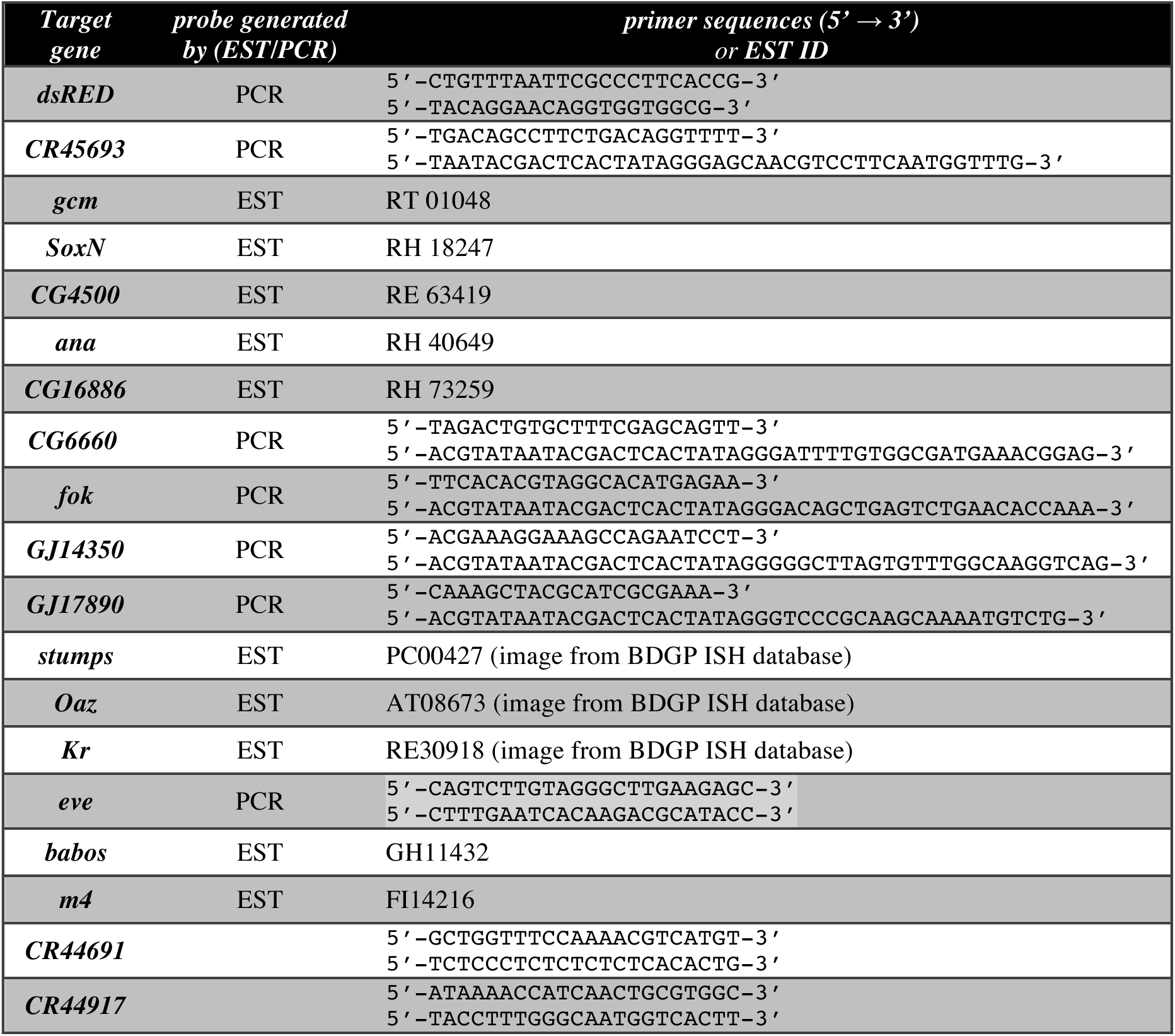
RNA in situ hybridization probe information is supplied. Probes were either generated from expressed sequence tags (DGRC clone identifiers listed), or from PCR products cloned into pCRII-Topo (PCR primers listed). 3 images corresponding to *stumps, Oaz,* and *Kr* ISHs were downloaded from the BDGP ISH database (*4*).

RNA in situ hybridizations were done according to standard procedures (*5*). Briefly, embryos of the genotypes used for cell isolation were collected for ca. 1.5hrs, aged for another 2.5hrs, then dechorionated and chemically cross-linked in 4% formaldehyde for 25 minutes before washing and dechorionation; fixed embryos were stored at -20°C until use. Antisense RNA in situ probes were generated by in vitro transcription using either ESTs obtained from the *Drosophila* Gene Collection (DGRC and BDGP), from PCR products using T7 priming sequences encoded in the reverse primers, or after T/A-cloning into pCRII (TOPO-Dual, Invitrogen). In all cases, inserts and orientations were sequence verified. Anti-sense probes were transcribed with DIG-UTP, hybridized overnight, and visualized using anti-DIG antibodies covalently linked to alkaline phosphatase in the presence of NBT+BCIP. Probes and primers are listed In Table S1. Standard imaging was done using Nomarski contrast on a Leica DMi8 microscope.

### Embryo collection and cell isolation for sequencing

For bulk sequencing, ~40 D. *mel.* or *D. vir.* stage 6 embryos were collected as described below. PBT was removed, embryos were shock frozen in liquid nitrogen and broken up using pestles. RNA was extracted by standard TRIZOL^®^ extraction. Quality of extracted RNA was assessed using a BioAnalyzer (Agilent Technologies, Wilmington, DE, USA). Libraries were generated from 200ng extracted RNA with the NEBNext^®^ Ultra^™^RNA Library Prep Kit for Illumina^®^ using poly(A) mRNA magnetic isolation. Samples were sequenced on the Illumina NextSeq500 platform using 75bp single end sequencing chemistry. Bulk sequencing libraries are deposited in GEO under accession numbers GSM 2494790 (*D. mel.,* mel_bulk) and GSM 2494791 (*D. vir.,* vir_bulk).

For single cell sequencing, *D. mel.* or *D. vir.* were allowed to lay eggs on apple juice-agar plates for 1 hr intervals and aged for ~2:30hrs (*D.mel*) or ~3:30hrs (*D.vir*) at room temperature. Embryos were dechorionated for 1 min in ~4% NaOCl and extensively washed with deionized water and rinsed with PBT (1×PBS, 0.1% Triton X100). Excess liquid was removed and embryos were transferred to 5% agarose gel slices stained light blue with Coomassie Brilliant Blue for contrast. Embryos were monitored under a stereomicroscope and stage 6 embryos were handpicked by morphological markers (beginning ventral invagination, start of germ band extension, no visible transverse furrow) and immediately transferred to ice-cold PBT (1×PBS, 0.1% Triton X100). Approximately 100 - 200 st.6 embryos were collected prior to dissociation.

Embryos were washed thoroughly using ice cold cell culture grade PBS to remove any residual detergent, resuspended in 1ml ice cold dissociation buffer (cell culture grade PBS, 0.01% molecular biology grade BSA) and dissociated in a dounce homogenizer (Wheaton #357544) with gentle, short strokes of the loose pestle on ice until all embryos were disrupted. The suspension was transferred into a 1.5 ml microfuge tube, cells were pelleted for 3’ at 1 000g at 4°C. The supernatant was exchanged for 1 ml fresh dissociation buffer. Cells were further dissociated using 20 gentle passes through a 22G × 2” needle mounted on a 5 ml syringe. The cell suspension was then gently passed through a 20 μm cell strainer (Merck, NY2002500) into a fresh 1.5ml microfuge tube, residual cells were washed from strainer using a small amount of dissociation buffer. Cells were pelleted again for 3’ at 1 000g at 4°C and resuspended in 100μl fresh dissociation buffer. The cell concentration was determined using a Neubauer counting chamber. Samples were fixed (*6*) by adding 4 volumes of ice-cold 100% methanol (final concentration of 80% methanol in PBS) and thoroughly mixing with a micropipette. Cells were stored at -20°C (up to several weeks without any noticeable decline in sequencing quality).

For Drop-SEQ runs, samples from 8 to 10 handpicking sessions were combined into batches. *D. mel.* and *D. vir.* samples were prepared separately and combined just before a Drop-seq run. For rehydration, cells were moved to 4°C and kept in the cold throughout the procedure. Cells were pelleted at 3 000g (1 000g for rep.1) for 5 minutes, resuspended in PBS + 0.01% BSA, centrifuged again, passed through a 35 μm cell strainer, counted and diluted for Drop-SEQ as described below. Single cell sequencing digital gene expression (DGE) matrixes are deposited in GEO under accession IDs GSM 2494783 – GSM 2494789 as indicated in Suppl. Table S2. An additional DGE of only the HQCs described in the paper (including their t-SNE cluster assignment) is available in GEO under accession ID GSE 95025.

**Suppl. Table S2:** Replicate run information for Drop-seq. CR_Hyd, number of cells recovered after rehydration; CR_CAF, number of cells recovered in combined aqueous flow.

| run ID | Drop-seq Run ID | Drop-seq Date | species mix | CR_Hyd | CR_CAF | final conc. (cells/µl) | GEO accession ID |
| --- | --- | --- | --- | --- | --- | --- | --- |
| rep.1 | ds015 | 2016-04-19 | yes (1:2) | 175 000 | 118 456 | 68 | GSM 2494783 |
| rep.2 | ds022_mix | 2016-07-05 | yes (2:1) | 100 000 | 72 360 | 40 | GSM 2494784 |
| rep.3 | ds022_mel | 2016-07-05 | no | 82 500 | 59 697 | 33 | GSM 2494785 |
| rep.4 | ds024a | 2016-09-02 | no | sample 1:<br>320 000 | 83 750 | 50 | GSM 2494786 |
| rep.5 | ds024b | 2016-09-02 | no |  | 83 750 | 50 | GSM 2494787 |
| rep.6 | ds024c | 2016-09-02 | no | sample 2:<br>212 500 | 58 625 | 35 | GSM 2494788 |
| rep.7 | ds024d | 2016-09-02 | no |  | 58 625 | 35 | GSM 2494789 |
| Total: |  |  |  | 890 000 | 535 263 |  |  |

### Drop-seq procedure, single cell library generation and sequencing

Monodisperse droplets of about 1 nl in size were generated using microfluidc PDMS devices (Drop-SEQ chips, FlowJEM, Toronto, Canada; either self-coated or pre-coated with Aquapel). Barcoded microparticles (Barcoded Beads SeqB; ChemGenes Corp., Wilmington, MA, USA) were prepared and flowed in using a self-build Drop-seq set up according to Macosko et al. 2015 (7) (Online-Dropseq-Protocol-v.- 3.1, http://mccarrolllab.com/dropseq/). Cells were loaded according to the table below in 1× PBS + 0.01% BSA. Droplets were collected in 50 ml Falcon tubes; typically, the collection tube was exchanged every ~12.5 minutes, corresponding to ~1 ml of combined aqueous flow volume (1 ml cells and 1 ml of beads). Droplets were broken promptly after collection and barcoded beads with captured transcriptomes were reverse transcribed, exonuclease-treated and further processed as described (*7*). The 1^st^ strand cDNA was amplified by equally distributing beads from one run to 48 PCR reactions (50 μl volume; 4 + 9 cycles). 10 μl fractions of each PCR reaction were pooled (total = 480 μl), then double-purified with 0.6x volumes of Agencourt AMPure XP beads (Beckman Coulter, A63881) and eluted in 12 μl 1 μl of the amplified cDNA libraries were quantified on a BioAnalyzer High Sensitivity Chip (Agilent). 600 pg cDNA library were fragmented and amplified (12 cycles) for sequencing with the Nextera XT v2 DNA sample preparation kit (Illumina) using custom primers enabling 3’- targeted amplification as described (*7*). The libraries were double-purified with 0.6x volumes of AMPure XP Beads, quantified and sequenced (paired end) on Illumina Nextseq500 sequencers (library concentration 1.8 p.; Nextseq 500/550 High Output v2 kit (75 cycles) in paired-end mode; read1 = 20 bp using the custom primer Read1CustSeqB (*7*), read 2 = 64 bp).

**Suppl. Table S3:**
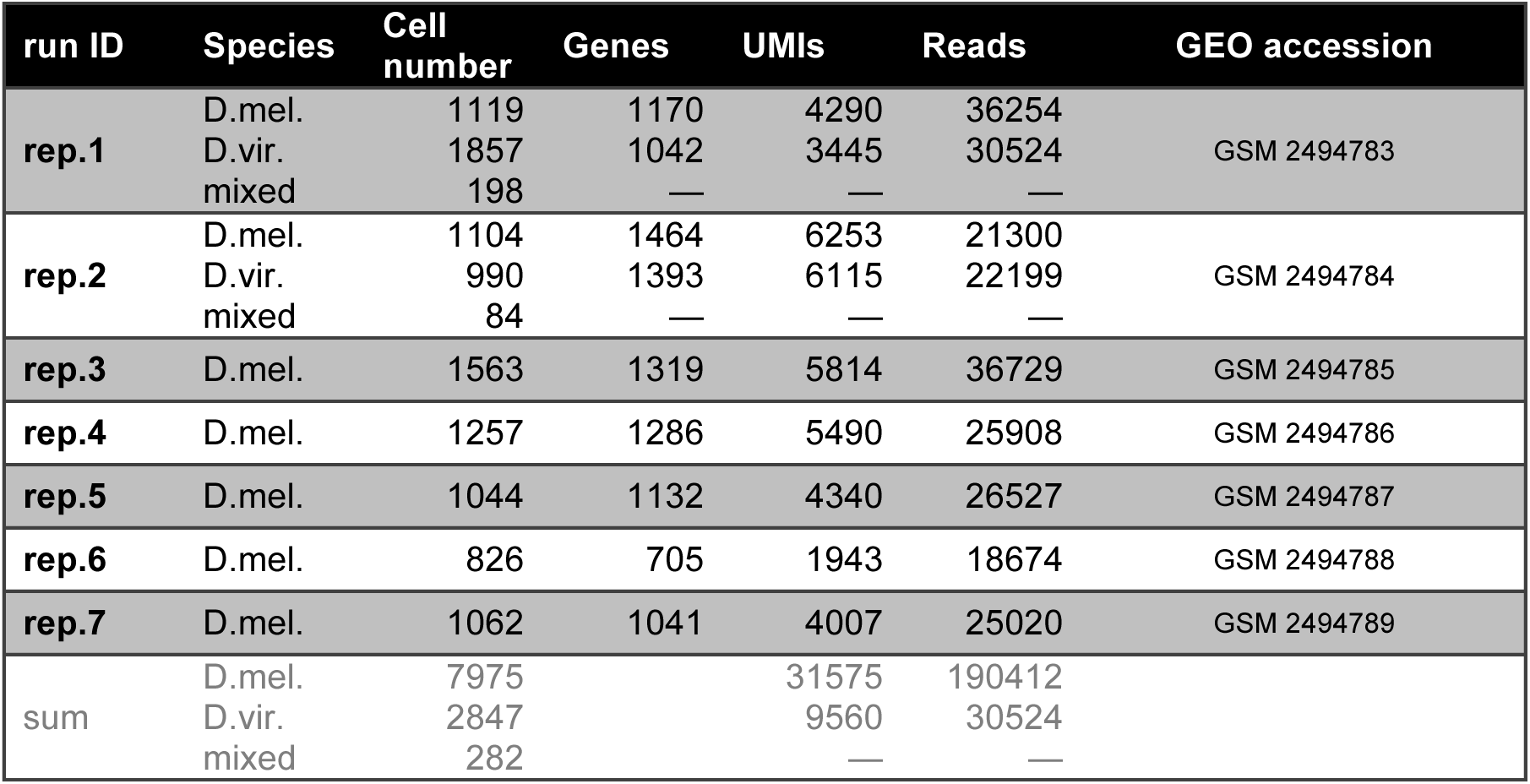
Drop-Seq statistics for stamps (cells) below the ‘knee’ and with >1000 UMIs. Genes, median number of genes/cell; UMIs, median number of UMIs/cell; Reads, median number of reads/cell.

| run ID | Species | Cell number | Genes | UMIs | Reads | GEO accession |
| --- | --- | --- | --- | --- | --- | --- |
| **rep.1** | D.mel. | 1119 | 1170 | 4290 | 36254 | GSM 2494783 |
|  | D.vir. | 1857 | 1042 | 3445 | 30524 |  |
|  | mixed | 198 | — | — | — |  |
| **rep.2** | D.mel. | 1104 | 1464 | 6253 | 21300 | GSM 2494784 |
|  | D.vir. | 990 | 1393 | 6115 | 22199 |  |
|  | mixed | 84 | — | — | — |  |
| **rep.3** | D.mel. | 1563 | 1319 | 5814 | 36729 | GSM 2494785 |
| **rep.4** | D.mel. | 1257 | 1286 | 5490 | 25908 | GSM 2494786 |
| **rep.5** | D.mel. | 1044 | 1132 | 4340 | 26527 | GSM 2494787 |
| **rep.6** | D.mel. | 826 | 705 | 1943 | 18674 | GSM 2494788 |
| **rep.7** | D.mel. | 1062 | 1041 | 4007 | 25020 | GSM 2494789 |
| sum | D.mel. | 7975 |  | 31575 | 190412 |  |
|  | D.vir. | 2847 |  | 9560 | 30524 |  |
|  | mixed | 282 |  | — | — |  |

### Single cell RNA-seq: data processing, alignment and gene quantification

The prepared libraries were sequenced in paired-end mode. We chose read 1 to be 20bp long, which sequences the cell barcode ‘stamp’ at positions 1-12 and the UMI at positions 13-20 (*7*), while avoiding reading into the poly(A) tail. The remaining 64 sequencing cycles were used for read 2. Sequencing quality was assessed by FastQC, while special attention was paid to the base qualities of read 1 to assure accurate cell and UMI calling. The last base of the UMIs generally showed a higher than expected T-content; we used the Drop-seq tools v. 1.12 to trim poly(A) stretches and potential SMART adapter contaminants from read 2, to add the cell and molecular barcodes to the sequences and to filter out barcodes with low quality bases. The reads were then aligned to a combined FASTA file of the *Drosophila* reference genomes (dm6; GCA_000005245) using STAR v. 2.4.0j with default parameters (*8*). Typically, around 85% of the reads were found to uniquely map to either of the species’ genomes; reads not uniquely mapping to a given genome were discarded. The Drop-seq toolkit was further used to add gene annotation tags to the aligned reads and to identify and correct bead synthesis errors, in particular base missing cases in the cell barcode. The Ensembl annotations BDGP6.86 and GCA_000005245.1.33 were used for *melanogaster* and *virilis* respectively. Cell numbers were estimated by plotting the cumulative fraction of reads per cell against the cell barcodes and calculating the inflection point. The DigitalExpression tool was used to obtain the digital gene expression matrix (DGE) for each sample.

### Species separation, cell filtering and data normalization

For efficient handling of mixed species samples and a first view of gene quantification, basic statistics and the doublet rates we used dropbead (https://github.com/rajewsky-lab/dropbead) (*6*). A threshold of 90% (UMIs mapping to one species) was selected to confidently declare reads corresponding to a single stamp as not representing a mixed-species doublet. *In silico* separation of the species under this threshold resulted in 7 individual DGEs for *D. mel.* and 2 for *D. vir.* (see Suppl. tables 2 and 3), which were subsequently pooled together by species, giving 7 975 and 2 847 cells, respectively. Within the pooled DGEs, cells containing more than 5 UMIs for mesodermal and dorsal ectodermal marker genes (see Suppl. Table 4) were removed as potential doublets. Pole cells were identified by containing at least 3 UMIs of *pgc* (*GJ22404* in *D. vir*) and were removed prior to clustering analysis and spatial mapping. Similarly yolk nuclei were identified by having at least 10 UMIs of yolk specific markers (see Suppl. Table 4). We further discarded cells expressing less than any 5 of the reference atlas genes used for mapping (threshold: 1 UMI), as our confidence in their mapping accuracy would be accordingly low. Genes mapping to the mitochondrial genome or with biotypes other than *protein coding, lncRNA* or *pseudogene* were also removed from the DGE. We normalized the UMI counts for every gene per cell by dividing its UMI count by the sum-total UMIs in that cell, and multiplying it by the number of UMIs that the deepest cell contained. Downstream analysis was performed in log space.

### Correlation of gene expression measurements

Correlations of gene expression levels between single-cell samples were computed by first sub-setting the DGEs of the two samples to the intersection of the genes captured in both libraries (typically ~10,000) and then computing the sum of gene counts across all cells in each library. Plotting of correlations is shown in log-space. For the correlation of our single-cell libraries against mRNA-seq, we converted the gene counts of the latter one into RPKMs and used the mean value of all isoforms for a given gene.

### Marker genes and germ layer assignment

We compiled sets of genes, which are well-known to be expressed specifically in the presumptive mesoderm (ME, ventral), neurogenic ectoderm (NE, lateral), or dorsal ectoderm (DE) (see Suppl. Table 4). Cells were then classified with respect to gross DV origin by computing a column-specific score per cell reflecting the average expression of any column's markers present in that cell. The scores were scaled and centered across every cell and subsequently across the three germ layers. Cells scoring higher than 1.3 in one of the columns were assigned to that column; otherwise they were designated ‘undetermined’.

**Suppl. Table S4:** *Drosophila* embryo germ layer marker genes (stage 5).

| Cell population | Genes |
| --- | --- |
| Dorsal ectoderm | *Ance, CG2162, Doc1, Doc2, egr, peb, tok, ush, zen* |
| Neurogenic ectoderm | *ac, brk, CG8312, l(1)sc, mfas, Ptp4E, sog, SoxN, vnd* |
| Mesoderm | *CG9005, Cyp310a1, GEFmeso, ltl, Mdr49, Mes2, NetA, ry, sna, stumps, twi, wgn, zfh1* |
| Yolk cells | *beat-IIIc, CG8129, CG8195, Corp, CNT1, sisA, ZnT77C* |
| Pole cells | *Pgc* |

### Identification of highly variable genes, PCA and t-SNE clustering

We used Seurat (*9*) for identifying the highly variable genes, i.e. genes with relatively high average expression and variability, in each of the pooled DGEs. Those sets were subsequently projected along the first few principal components to encompass more genes (see Satija et al., 2015 for more details on this procedure). Principal component analysis was performed over these extended sets. The first 16 or 18 principal components were identified as statistically significant and were used as input for t-SNE clustering of *D. mel* or *D. vir.* cells, respectively. We used Seurat to identify the marker genes for each of the clusters in the t-SNE representation.

**Suppl. Table S5:**
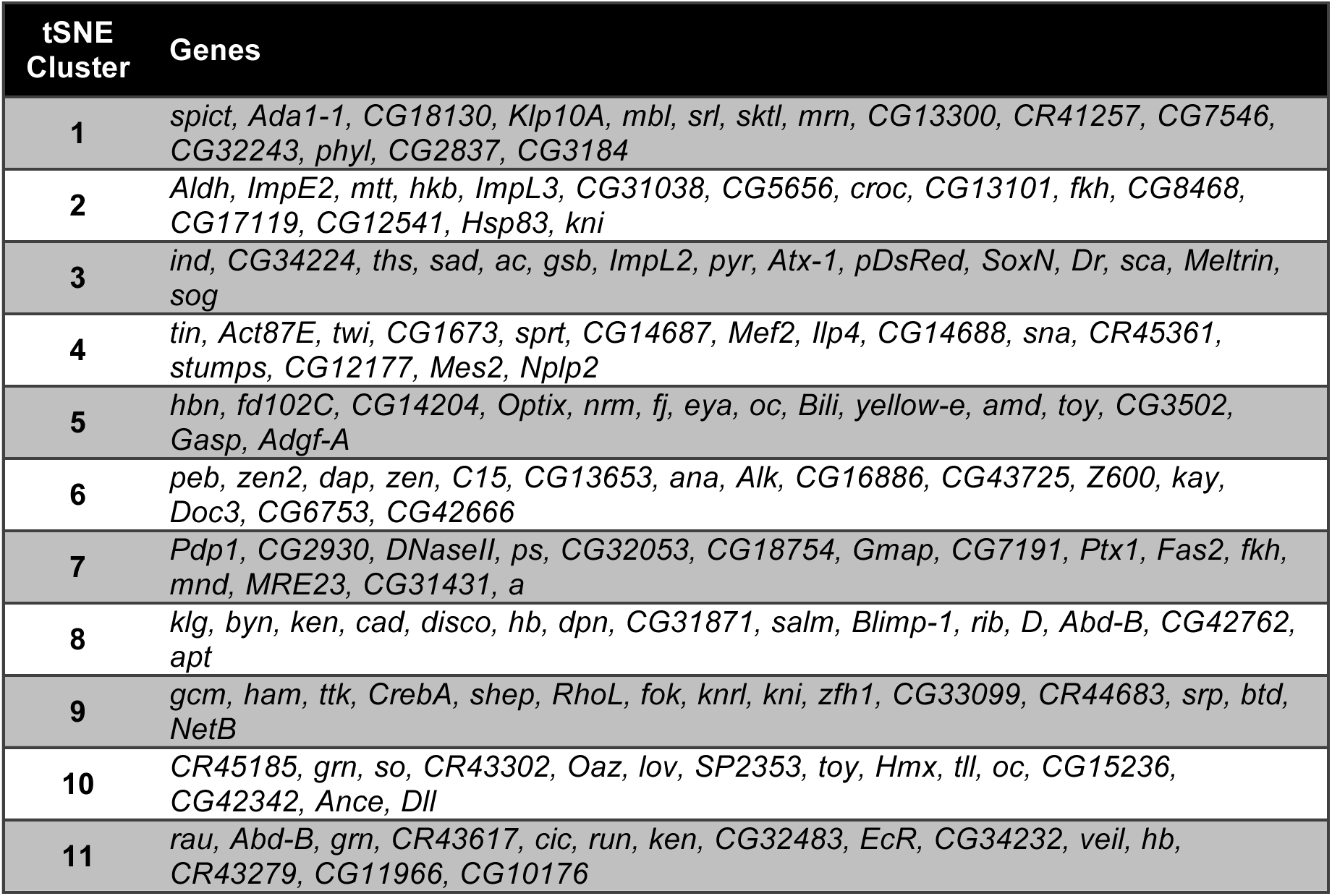
Highly variable genes in tSNE clusters.

### Binarization of the BDTNP reference atlas

The BDTNP reference atlas (*10*) quantifies the relative mRNA levels of 84 genes in the context of a virtual stage 5 *D. mel* embryo at pseudo-nuclear resolution. We selected the latest of the 6 time points, i.e. stage 5 with ~100% cellularization, for mapping, as this is the closest match to our data – the earliest we were able collect embryos was beginning developmental stage 6, because cellularization in the *Drosophila* embryo does not complete until the end of stage 5, which means our embryos are developmentally within ~10 minutes of the high resolution BDTNP marker gene atlas used. Due to bilateral symmetry only half of the embryo is relevant for spatial mapping; hence we selected the first 3 039 nuclei. We converted the continuous gene expression levels into binarized on/off states that would guide spatial mapping. We chose a binarization threshold for each gene individually, by inspecting a range of thresholding values and comparing the resulting pattern with published in situ hybridizations (*4, 11*). In cases where the BDTNP atlas thresholding resulted in clear discrepancy with published in situ data, the reported expression was given primacy and the atlas was manually adjusted. Out of 3 039 locations of the gene atlas, 2 937 exhibit unique gene combination patterns upon binarization. The final position dependent expression matrix of the binarized in situs used for spatial mapping is included in Suppl. Table 5.

### Binarization of Drop-seq data

We binarized the gene expression levels for the 84 genes that guided the mapping as follows. First, we computed the gene correlation matrix of the 84 genes with the binarized version of the BDTNP atlas. Next, for a given gene and only considering the Drop-seq cells expressing it we computed a quantile value above (below) which the gene would be designated ON (OFF). We sampled a series of quantile values and each time the gene correlation matrix based on this binarized version of our data versus the binarized BDTNP atlas was computed and compared by calculating the mean square root error between the elements of the lower triangular matrices. Eventually, we selected the quantile value 0.23, as it was found to minimize the distance between the two correlation matrices.

### Spatial mapping algorithm

Having binarized the BDTNP atlas and our data as described above, we aimed at assigning a probability score for each of our cells to each of the 3 039 bins of the database as follows. We compared a given cell to all bins by counting the number of genes found ON (OFF) in both the cell and a given bin, as well as the number of genes which disagreed between the cell and the given bin. This led to a calculation of 3 039 confusion matrices for each cell. In a given cell-bin combination, we interpreted the OFF-OFF cases as true negatives (tn), the ON-OFF cases as false positives (fp), the OFF-ON cases as false negatives (fn) and the ON-ON cases as true positives (tp). We employed the Mathews correlation coefficient (MCC) to weight the confusion matrices and assign a cell-bin score. The MCC scores were subsequently exponentiated.

### Simulations to assess bin coverage & cell mapping confidence

In general a given cell exhibited high scores with more than one positional bins (see also heat map of MCC scores in Fig 4A). We performed simulations to assess whether the obtained high scores could have been generated by chance. First, we permuted the values of the 84 genes for every cell of our data 100 times, generating thus 129 700 simulated cells. We mapped the simulated cells onto the reference atlas via the algorithm described above and calculated then the maximum MCC scores per bin. Their distribution was significantly lower than the corresponding distribution arising from the mapping of real cells (data not shown). Particularly, more than 87% of the bins had at least one MCC score with a p-value smaller than 0.05, leading to the conclusion that we effectively covered more than 87% of the embryo with high confidence (Fig. 4C). Similarly, we permuted the 84 genes 40 times for every bin of the reference atlas, generating thus 121 560 simulated bins, upon which we mapped our Drop-seq data. Comparing the distribution of the maximum MCC scores with the one corresponding to the real bins, we assessed that 878 cells were assigned to the embryo with very high confidence (p-value < 0.05, data not shown).

### Computation of gene expression patterns

For the majority of the cells, several positional bins scored highly. Therefore, we adopted a probabilistic mapping strategy, instead of fixing every cell to only its highest probability location. For a given gene, we multiply the vector of its normalized expression across cells with the MCC scores per bin. This multiplication is element-wise and penalizes cases for which the gene is either very lowly expressed (even though the scores for that cell could be very high) or the score for a bin is very low (indicating that the cell expresses the gene but maps better to another bin). We sum the products row-wise, obtaining thus a number for each bin. We then binarize the vector of normalized gene expression and repeat the above process, resulting in a second 3309 dimensional vector. We divide the first vector by the second, element-wise, to account for the different number of cells that could express the gene. The resulting vector, q, is further normalized through q/(1+q) to reduce outlier effects. A threshold is then imposed that sets the values of q to zero that lie below the quantile corresponding to that threshold.

### Plotting of gene expression patterns

We used the coordinate system of the sixth time point of the BDTNP database to depict our computed expression patterns. As the embryo is bilaterally symmetric and we mapped our cells to the first 3 039 bins, we mirrored the final scores for the remaining 3 039 bins across the A/P-D/V plane to obtain a complete picture of the embryo. For the lateral view, we display the first 3 039 bins plotted in x and z. For the ventral and dorsal views, the bins with z<0 or z>0 were selected and their x/y coordinates were plotted. For the anterior and posterior view, the bins with x<0 or x>0 were selected and their y/z coordinates were plotted. The red to grey color scale shows high to low expression per positional bin.

### Noise simulations to assess vISH threshold values

We performed simulations in order to identify meaningful threshold values when computing a vISH. Using the raw data of the 1 297 cells, we generated for every gene and for every cell a random Poisson number, with a mean and variance equal to the number of UMIs for that cell and gene; we repeated this process 50 times. We verified that the correlations of gene expression levels were high across the samples (Pearson R > 0.97, data not shown), but the cell-to-cell correlations were much lower (Pearson R ~ 0.7, data not shown). We spatially mapped these 50 simulated samples. For a given gene we computed a vISH for thresholds in the range [0.5, 0.98] with steps of 0.01. We then computed the per bin standard deviations across the 50 calculated vISH and their sum. As higher thresholds imply fewer bins, this sum is expected to minimize for threshold=0.98. However, for some genes, especially the well-expressed ones, the distribution of these sums against the threshold values showed local minima, which in some cases were even global (Sup. Fig. 6A), revealing robustness of these threshold values against Poisson noise simulated data. Visualizing the vISHs with these obtained values and comparing the patterns with in vivo spatial expression data (*4, 11*) showed good agreement. For the lowly expressed genes our simulations could not provide useful insights on threshold values, even when modeling noise with a negative binomial distribution, allowing for different variances.

### Clustering and discovery of archetypal in situ patterns

Having computed a set of gene expression patterns, we computed their distance matrix in order to discover gene expression archetypes. The gene set was selected as the union of the highly variable genes, the three column markers and the genes used for the spatial mapping. Agglomerative clustering was performed to identify the number of parent clusters. We averaged the expression patterns of the genes belonging to each of the parent clusters, thus producing 10 representative in situs of the corresponding identified archetypal classes (Sup. Fig. 7B).

### Discovery of new markers with localized expression patterns

We restricted our survey for new markers of expression patterns to genes either not reported in, or annotated as ‘no staining’ in the BDGP *in situ* database. We computed the expression patterns for this large set of genes and visually inspected these virtual *in situ* hybridizations. For validation, we chose genes with distinct localized expression patterns.

### Correlation of Drop-Seq and nCounter gene expression analysis

To assess the quantitativeness of the Drop-Seq data gene expression data with respect to NanoString nCounter data from individual stage-matched embryos (beginning gastrulation) (*12*), the DGE was subsetted to the genes included in the nCounter analysis. For each gene a total count across all cells was calculated. The gene *Fdy* was excluded because it resides on the Y-chromosome and the nCounter value based on 3 embryos cannot be representative of a random male/female mixture of cells. For genes which were counted using two primer pairs in the nCounter experiment (genes) the higher value was used in each case, assuming that this reflects expression values obtained by poly-A selection more closely due to inclusion of more transcript isoforms. For each gene, the total DGE counts and the total nCounter (gastrulation stage) counts were converted to relative expressions by dividing each gene count by the sum of all genes in the respective dataset. Correlation plot is shown in log-space.

### Gene Ontology Term Enrichment analysis

For the GO term analysis the Comprehensive R Archive Network (CRAN) package “gProfileR” was used. All genes detected in the Drop-seq experiment were used as the background model. For the cluster-specific GO term enrichments, a list of genes identified as significantly overrepresented in a cluster was assembled for each cluster using the “FindAllMarkers” function from Seurat (*9*) and requiring an average difference (avg_diff) > 0 and a p-value (p_val) < 0.05. Cluster specific enrichment of terms is displayed on a blue scale where p-value > 0.05 and on a red scale were p-value < 0.05 (Suppl. Fig. S3 and S4). P-values are displayed as -log10 (p-value). Rows are clustered by complete linkage clustering. The inclusive heatmap for biological process terms (Suppl. Fig. S4) was subsetted to exclude less meaningful terms and is shown in Suppl. Fig. S3.

## Supplemental Figures and Figure Legends

**Suppl. Figure S1:**
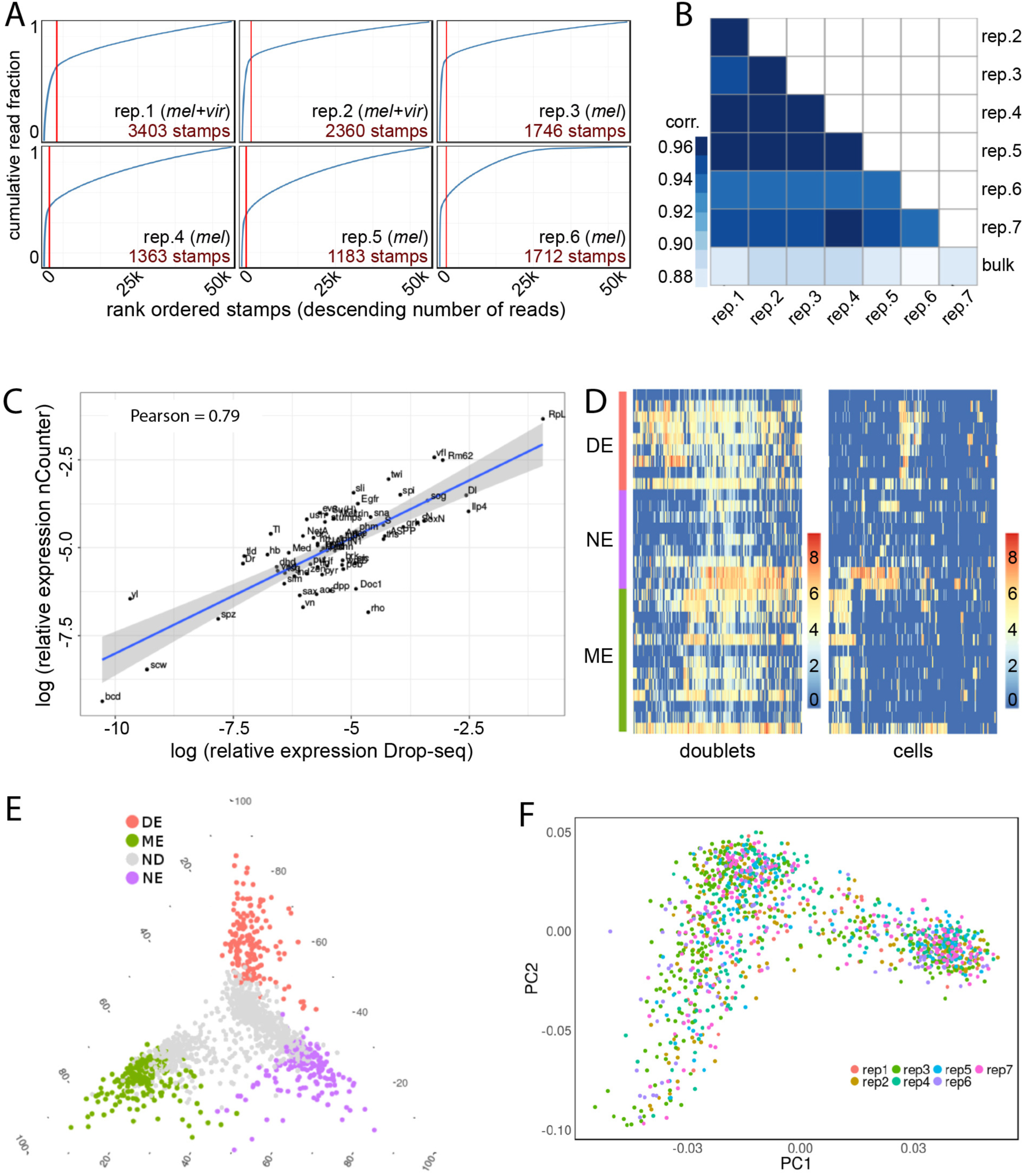
Drop-seq data quality and reproducibility. (A) Knee plots. Cumulative fraction of reads vs. cells rank ordered by decreasing read number are shown for 6 separate Drop-seq runs. The inflection point (red line) marks the number of recovered cells. (B) Correlation matrix comparing aggregated UMIs in Drop-seq replicates and whole embryo mRNA sequencing (bulk) for *D. mel.* (C) Correlation of relative gene expression between aggregated UMIs in Drop-seq and nCounter measurements (*12*). (D) Doublet identification and exclusion. Heat maps of cells (columns) clustered by expression of DE, NE and ME marker genes (rows). Doublets (left) show expression of several marker classes whereas a random subsample of single cells (right, doublets excluded) shows much more exclusive marker class expression. Coloring indicates normalized expression levels (log2(ATPM+1)). (E) Ternary plot indicating scores for marker expression. Grey cells could not be unambiguously assigned to any of the ME, NE, or DE populations. (F) Absence of batch effects. PCA of 1 297 mapped *melanogaster* cells; cells are colored by Drop-seq runs.

**Suppl. Figure S2:**
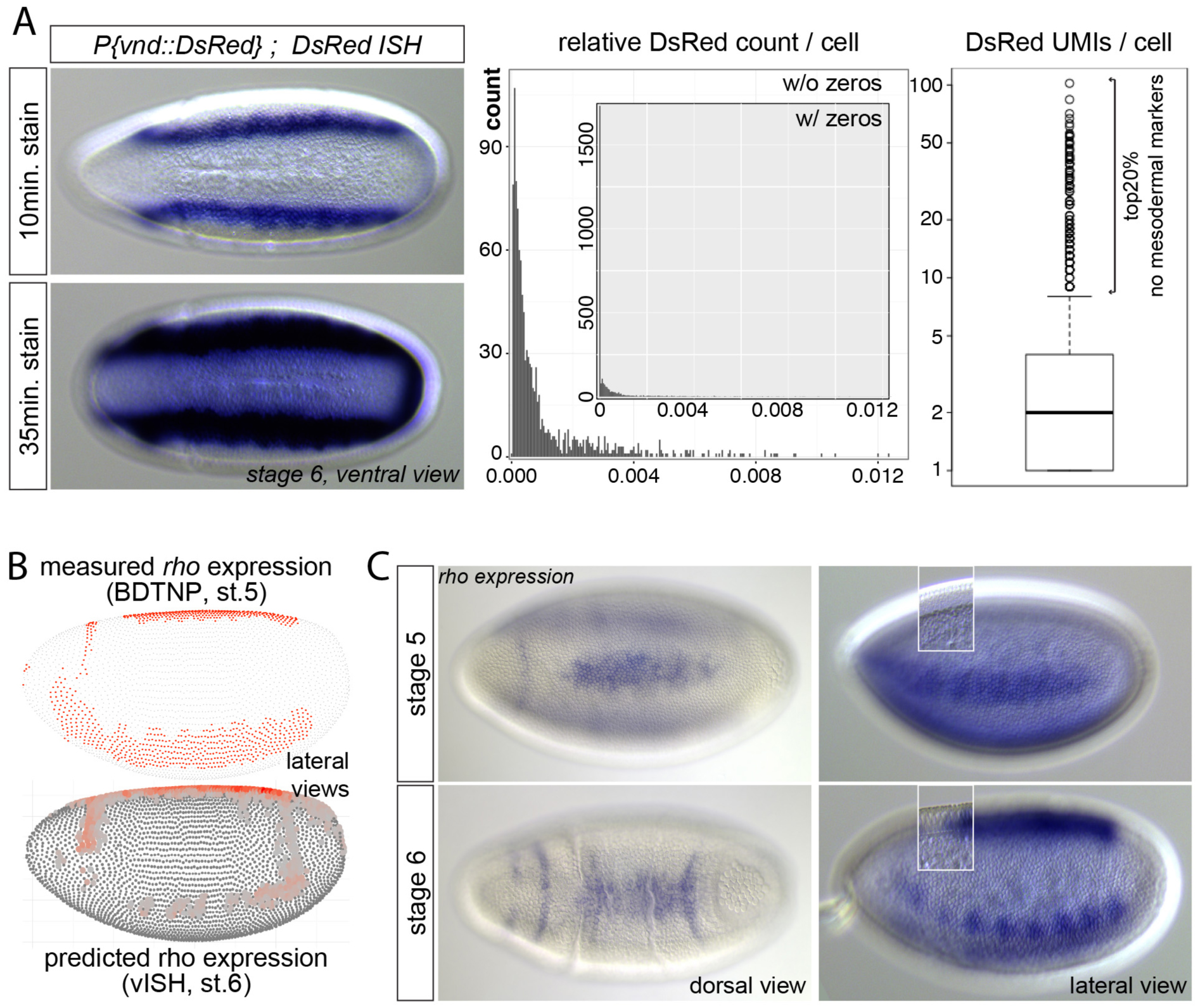
Spatial expression of *DsRed* and rho *in vivo*. (A) Left – The *vnd::DsRed* transgene is expressed primarily in the ventral neurogenic ectoderm in 2 lateral stripes, but excessive staining shows weaker expression also in the presumptive mesoderm. Middle – Distribution of *DsRed* reads among sequenced cells; relative *DsRed* UMI count per cell among ~2 000 sequenced cells, excluding or including (inset) cells that show zero *DsRed* UMIs. Right – boxplot of *DsRed* UMI counts across cells; cells expressing *DsRed* highly do not express mesodermal markers. (B) Measured rho expression (top) and vISH predicted rho expression disagrees, especially in ventro-lateral regions. (C) RNA ISH against rho shows continuous lateral stripes at stage 5 (top-right, see inset for cellularization state, focal plane adjusted), but patchy expression at stage 5/6 (bottom), similar to vISH (compare panel B, bottom). Dorsal expression (left) does not significantly change, measured and prediction agree. Embryos in the left panels were stained for a shorter amount of time indicating higher dorsal expression, which also agrees with the vISH prediction. Embryos are oriented anterior left, DV rotation as indicated.

**Suppl. Figure S3:**
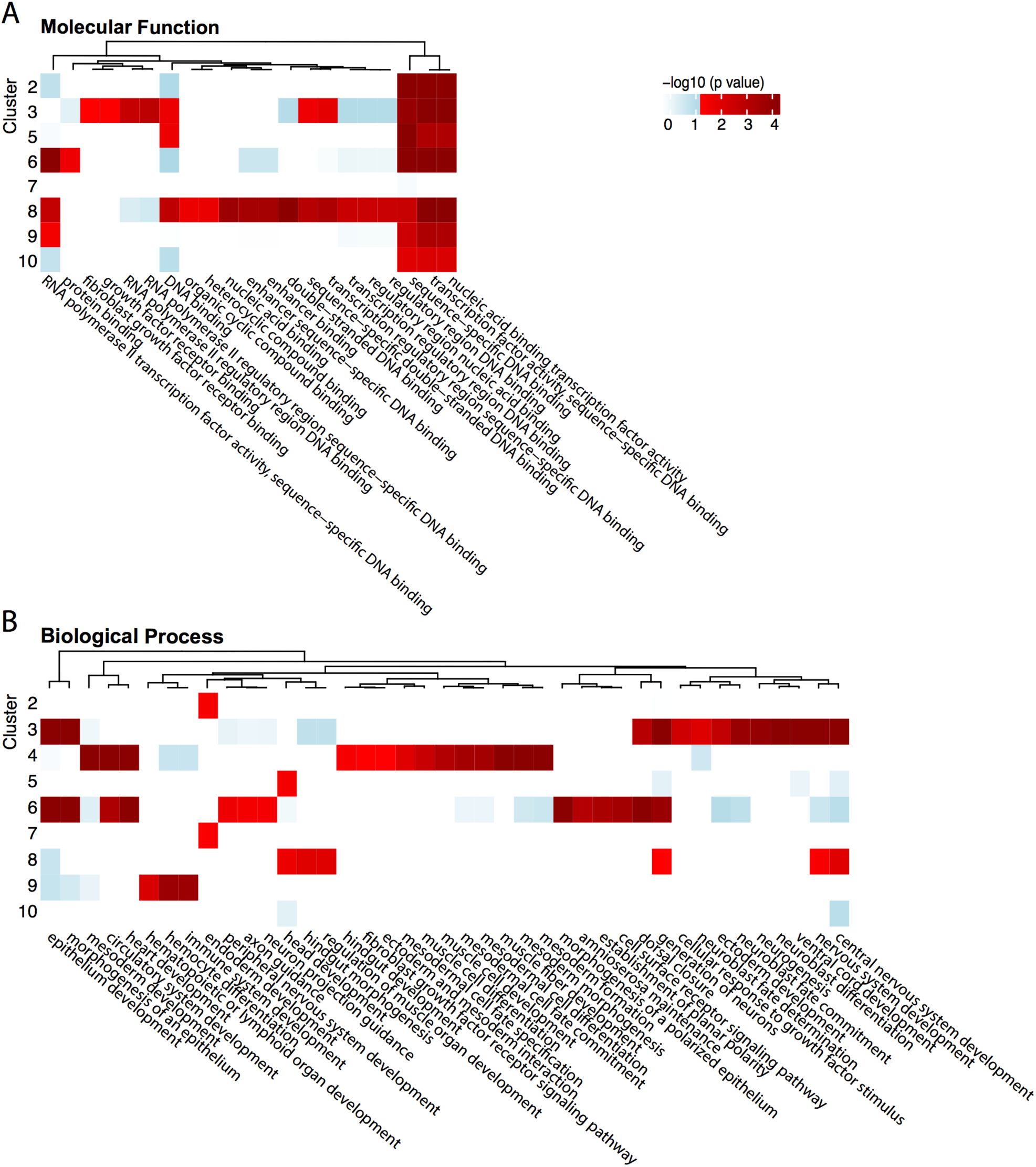
GO term enrichment across single cell t-SNE clusters. GO term enrichment indicates distinct molecular and biological functions associated with different t-SNE clusters. Molecular Function (A) and Biological Process (B) GO term enrichment among t-SNE cluster-enriched genes. Cluster numbers correspond to t-SNE clusters as indicated in Fig. 3C. Color Scale indicates adjusted p-value. The heat map in (B) has been subset for clarity; the complete map is available in Suppl. Fig. S4.

**Suppl. Figure S4:**
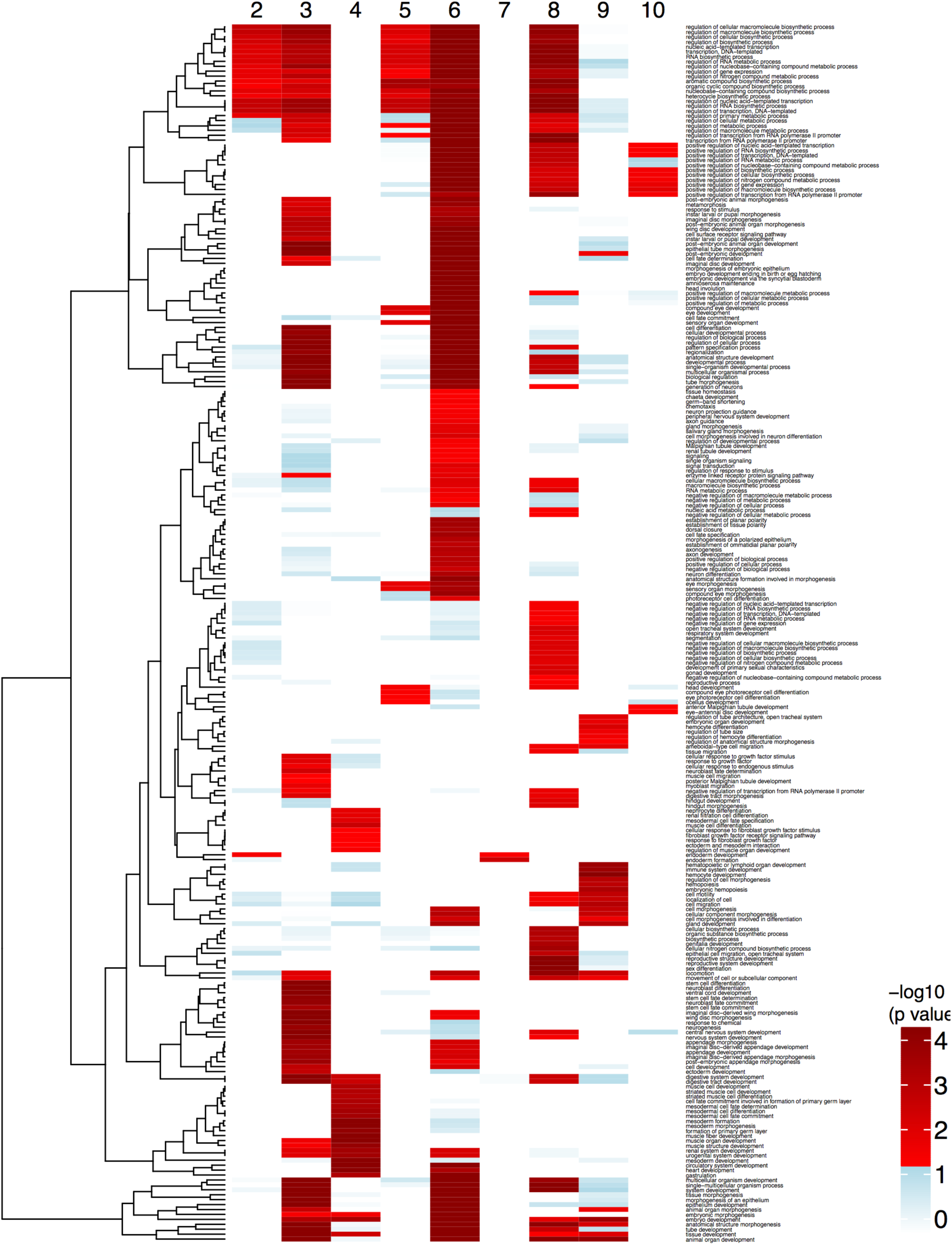
Biological Process GO term enrichment across single cell t-SNE clusters (inclusive map corresponding to Sup. Fig. S3B). An inclusive map of Biological Process GO term enrichment among t-SNE cluster-enriched genes. Cluster numbers correspond to t-SNE clusters as indicated in Figure 3C. Color Scale indicates adjusted p-value.

**Suppl. Figure S5:**
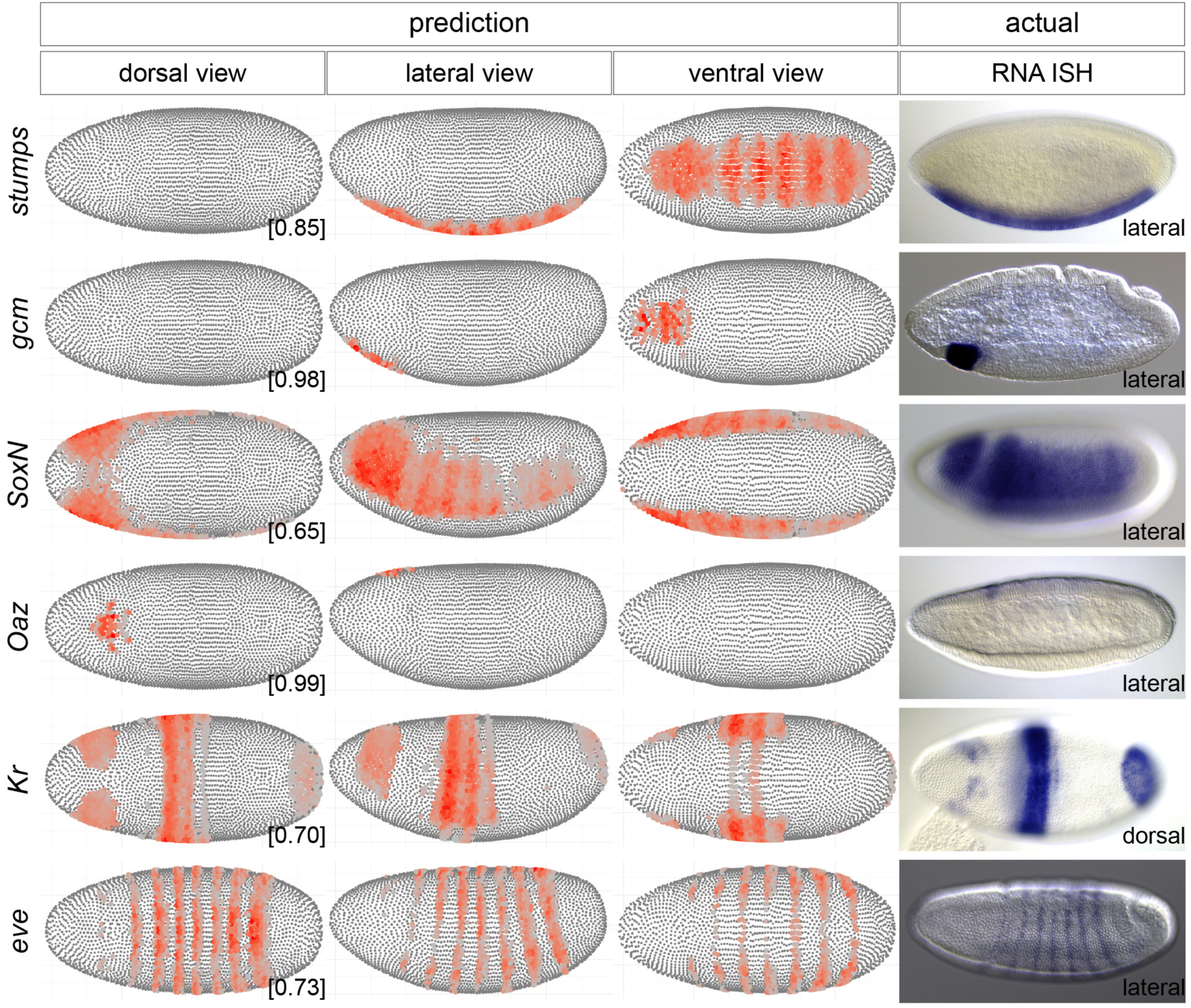
vISH predictions and experimental validations for known marker genes. vISH predictions of known marker genes are depicted anterior left in three DV rotations as indicated. Actual *in vivo* expression shown by RNA *in situ* hybridization (right), Embryo orientation is anterior left, DV rotation as indicated, shown are developmental stage 5 or 6. (*stumps, Oaz* and *Kr* images from BDGP.)

**Suppl. Figure S6:**
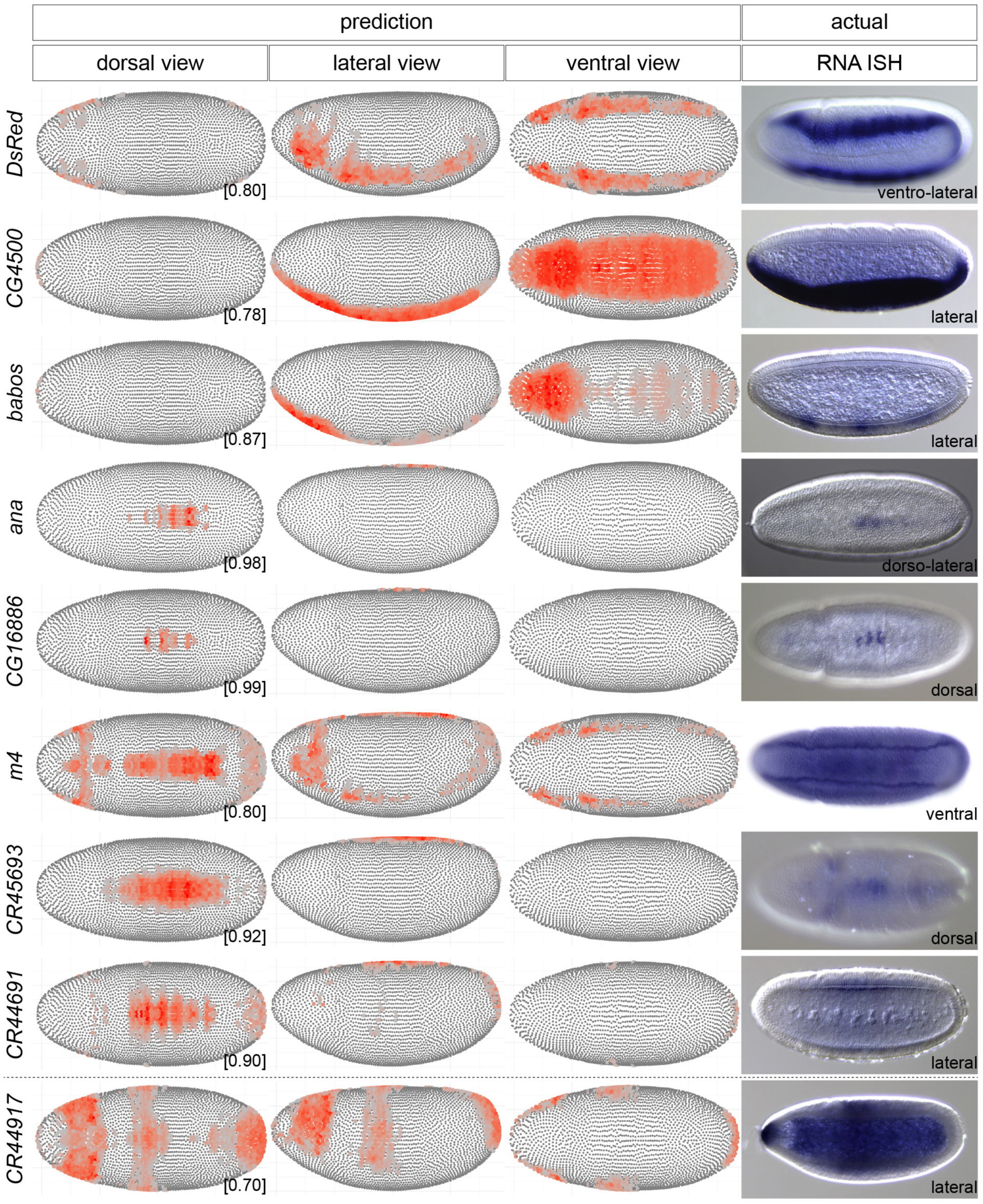
vISH predictions and experimental validations for novel marker genes, including long non-coding RNAs. *De novo* spatial expression predictions by vISH (left) depicted anterior left in three DV rotations as indicated. Actual *in vivo* expression validated by RNA *in situ* hybridization (right), Embryo orientation is anterior left, DV rotation as indicated, shown are developmental stage 5 or 6.

**Suppl. Figure S7:**
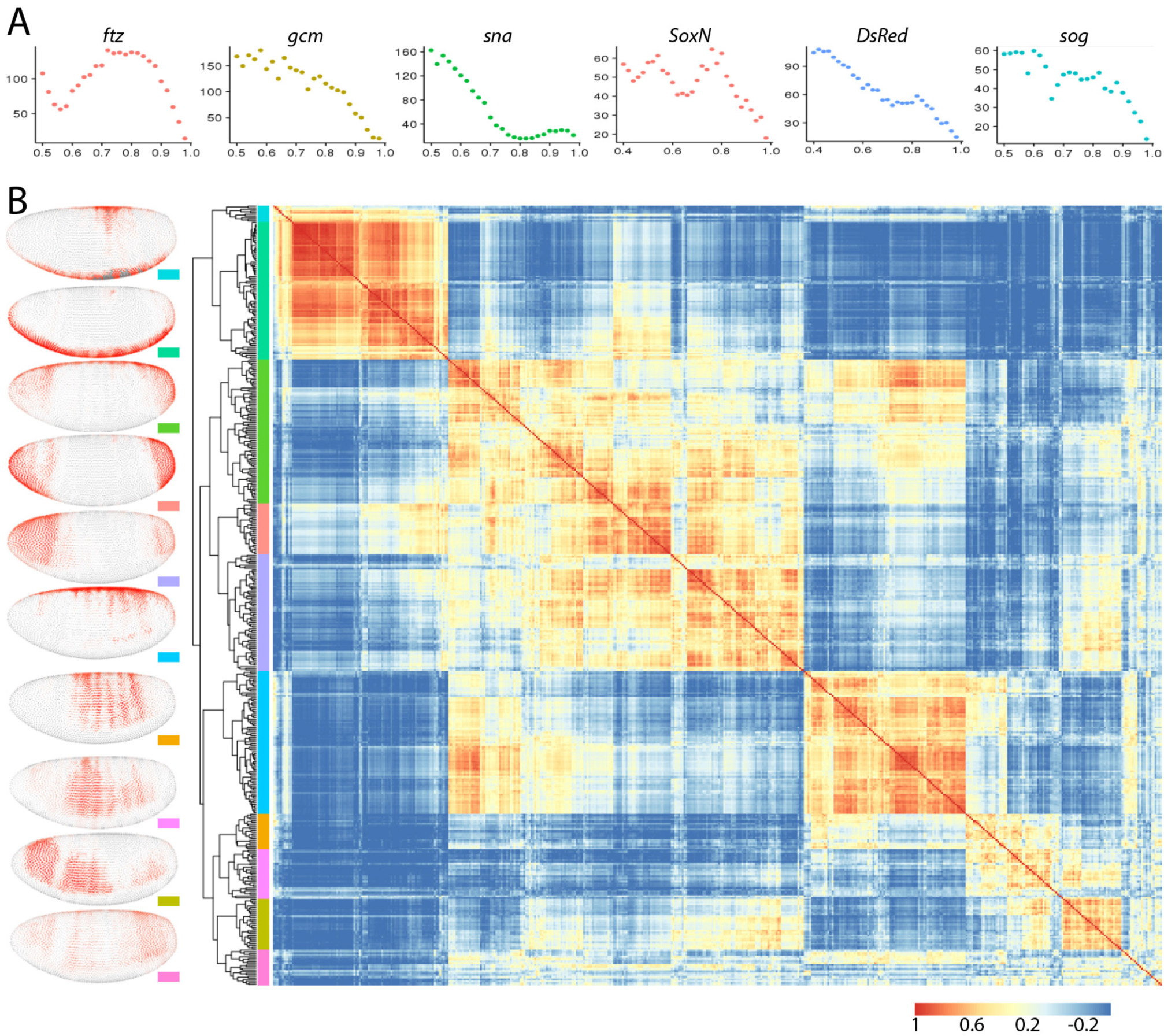
vISH thresholds and discovery of gene expression archetypes. (A) Assessment of threshold values via Poisson noise simulations (see also Suppl. Materials and Methods for details). Thresholds (x-axis) are plotted against the sum of standard deviations across all bins (y-axis). As more stringent thresholds imply less bins, the above sum is expected to decrease monotonically. The existence of additional local minima in the cases of *ftz, sna, SoxN* and *sog* reveals threshold values around which vISHs are meaningful. (B) The 476 most highly varied genes were clustered according to embryo-wide expression similarity. Heat map shows clustering of the correlation matrices of the computed vISHs with a global threshold value equal to 0.75. Average expression of genes within parent clusters reveals archetypal patterns shown as vISHs to the left. Color scale indicates correlation of Euclidean distances.

